# *Trichomonas vaginalis* targets *Lactobacillus jensenii* via pseudopodia-independent phagocytosis and secreted lysozyme TvGH25

**DOI:** 10.64898/2026.07.07.735988

**Authors:** Nadine Zimmann, Michal Havelka, Alois Zdrha, Jana Procházková, Tamara Smutná, Petr Rada, Zdenek Verner, Adam Hart, Jaya Sharma, Jacob Biboy, Daniela Vollmer, Waldemar Vollmer, Jan Tachezy

## Abstract

A low abundance or absence of protective lactobacilli during acute trichomoniasis is a well-known phenomenon that is associated with *T. vaginalis* (TV) infection. However, a crucial question that remains unanswered is whether alterations in the lactobacilli population precede TV infection or whether the parasite plays an active role in lactobacilli disappearance. Our findings showed that TV efficiently phagocytosed one of the dominant *Lactobacillus* species *L. jensenii* (LJ). Phagocytosis proceeds via a pseudopodia-independent mechanism reminiscent of sinking with a preference for viable cells. The presence of viable LJ leads to an increase in secretion of 27 TV proteins, including TvGH25 lysozyme. This enzyme cleaves peptidoglycan, a major component of the bacterial cell wall. TV overexpressing TvGH25 effectively lowers the bacterial cell count, evidencing the enzyme’s antimicrobial potential. These data support the notion that TV cells can suppress the *Lactobacillus* population through a combination of targeted secretory response and phagocytic activity, revealing novel potential targets for developing alternative therapeutic strategies against trichomoniasis.

**Significance Statement:** *Trichomonas vaginalis* (TV) is a sexually transmitted parasite that causes trichomoniasis and is connected to the disruption of the healthy vaginal microbiome, dominated by *Lactobacillus* species. However, the nature of the interactions between TV and *Lactobacillus* is poorly understood. In this study, we show that TV uses an unusual form of pseudopodia-independent phagocytosis to engulf *L. jensenii* alongside a targeted secretory response to the bacterial encounter, involving TvGH25 lysozyme. We found that TV acquired this enzyme by lateral gene transfer from bacteria and repurposed it against bacteria to degrade the major bacterial cell wall component peptidoglycan. TvGH25 thus represents an effective component of TV’s antibacterial arsenal. Our findings provide new insights into the mechanistic disruption of the protective *Lactobacillus* microbiota.

## Introduction

*Trichomonas vaginalis* (TV) is a flagellated parasitic protist responsible for the most prevalent non-viral sexually transmitted infection, with 156 million new cases reported annually (1). In women, the infection can lead to vaginitis, and is associated with an increased risk of HIV transmission, preterm birth, low birth weight, and cervical cancer. In contrast, most infected men remain asymptomatic; however, chronic infections have been linked to an elevated risk of prostate cancer (2–5).

Within the vaginal mucosa, TV actively phagocytoses a variety of host cells, including epithelial cells, lymphocytes, erythrocytes, as well as cellular debris and vaginal microorganisms like yeast and bacteria (6–9). In addition to internalization, TV secretes numerous biologically active proteins, such as adhesins that promote cytoadhesion and phagocytosis, and a variety of proteases that degrade nutrients, microbial cell walls, and host cell structures (4, 10, 11). Thereby, TV not only gains access to valuable nutrients but also overcomes host defense mechanisms. A healthy vaginal microbiome is typically dominated by Lactobacillus species (12), which help prevent the overgrowth of pathogenic microbes and support the immune defenses within the vagina; however, a TV infection is associated with a disruption of the healthy microbiome and an imbalance in the microbial community (12, 13). Dysbiotic microbiota can, in turn, exacerbate symptoms such as inflammation and promote TV colonization, which may result in chronic TV infections. In women, the vaginal bacterial community is grouped in five community state types (CSTs), four of which predominantly consist of *Lactobacillus* species, i.e., *L. crispatus* (CST-I), *L. gasseri* (CST-II), *L. iners* (CST-III), and *L. jensenii* (LJ, CST-V) (14). CST-IV does not have a dominant *Lactobacillus* species and is characterized by a higher proportion of strict anaerobes such as *Prevotella*, *Atopobium*, and *Gardnerella* (14). The CSTs all differ in their protective abilities towards pathogens, with CST-I and CST-II reportedly being the most protective (15–20). Specifically, *L. gasseri* was shown to inhibit TV adhesion to ectocervical cells (17, 18), while dysbiotic bacteria stimulate the parasite adhesion (21). LJ (CST-V) has been shown to generally benefit vaginal health (22, 23) and pregnancy outcomes (24, 25). In contrast, *L. iners* (CST-III) is considered a transitional species with an ambiguous role in the vaginal community: it coexists with both beneficial and pathogenic microbiota, indicating it provides limited protective effects (20, 26, 27), whereas CST-IV has been associated with an 8-fold higher risk of a TV infection (12, 14, 20, 28).

Intriguingly, it is not yet known whether and to what extent TV actively causes alterations in the healthy microbiome. To contribute to the elucidation of this critical question, we investigated interactions between TV and LJ on the cellular as well as protein level. Initially, we tested TV phagocytic activity against LJ, and then continued with further detailed investigations of phagocytosis and TV response to this organism. We found that LJ cells stimulated the TV secretion of biologically active proteins, including TvGH25 lysozyme. This enzyme of bacterial origin cleaves bacterial peptidoglycan and displays antimicrobial potential. TvGH25 thus represents a new potent component of TV’s antibacterial machinery.

## Results

### *T. vaginalis* is highly phagocytic to *L. jensenii*

To evaluate the phagocytic activity of TV to the CST-V dominant protective *Lactobacillus* species LJ, TV and LJ were co-incubated in a 1:10 ratio for 5, 15, and 30 min at 37°C, and then phagocytosis was evaluated. Initially, we used a microscopy assay that allowed us to distinguish engulfed lactobacilli from those attached to the TV surface (Fig. 1A). The microscopy assay revealed a time-dependent increase in TV cells with engulfed bacteria (Fig. 1B). Phagocytosis was observed as early as 5 min of co-incubation, when about 10% of TV cells were positive for LJ. After 30 min, about 30% of TV cells contained LJ (Fig. 1B). However, in the microscopy assay, only a limited number of cells can be evaluated. Thus, next, we investigated the phagocytic activity by imaging flow cytometry followed by a downstream machine-learning pipeline to distinguish between surface-attached LJ and phagocytosis. This approach enabled us to analyze thousands of TV cells while ensuring accuracy. Therefore, TV was co-incubated with BacLight™ Red-stained lactobacilli for up to 30 min at 37°C, and as a control, the cells were incubated at 15°C, at which TV phagocytic activity is significantly reduced (Fig. 2). The imaging flow cytometry analysis revealed four groups based on the BacLight Red signal (Fig. 2A, 2B): (i) Phagocytosis, with a sharp internalized LJ signal; (ii) Adjacent, with LJ signal on the cell surface or in proximity to TV’s; (iii) Homogenous, with BacLight Red signal homogenously distributed across the TV cell; (iv) No BacLight Red signal. The automated image analysis confirmed high phagocytic activity to LJ, with an average of 30% of TV cells having ingested LJ after 30 min of co-incubation (Fig. 2C). While the phagocytosis group grew significantly over time, the adjacent group decreased in 30 min. TV cells with a homogenous BacLight Red signal (approximately 2% in 30 min) most likely represent released stain from digested bacteria (Fig. 2D). Next, we tested the phagocytic activity of TV to heat-inactivated LJ (HiLJ). Therefore, TV was co-incubated with either BacLight™ Red-stained HiLJ or viable LJ (vLJ) for 30 min at 37°C, followed by imaging flow cytometry analysis. The co-incubations revealed a phagocytic preference for viable LJ. About 37% of TV had phagocytosed vLJ after 30 min, while phagocytosis of HiLJ averaged at 23% (Fig. 2E). These results suggest that phagocytosis of LJ is supported by but does not depend on native components actively produced by LJ.

**Figure 1.**
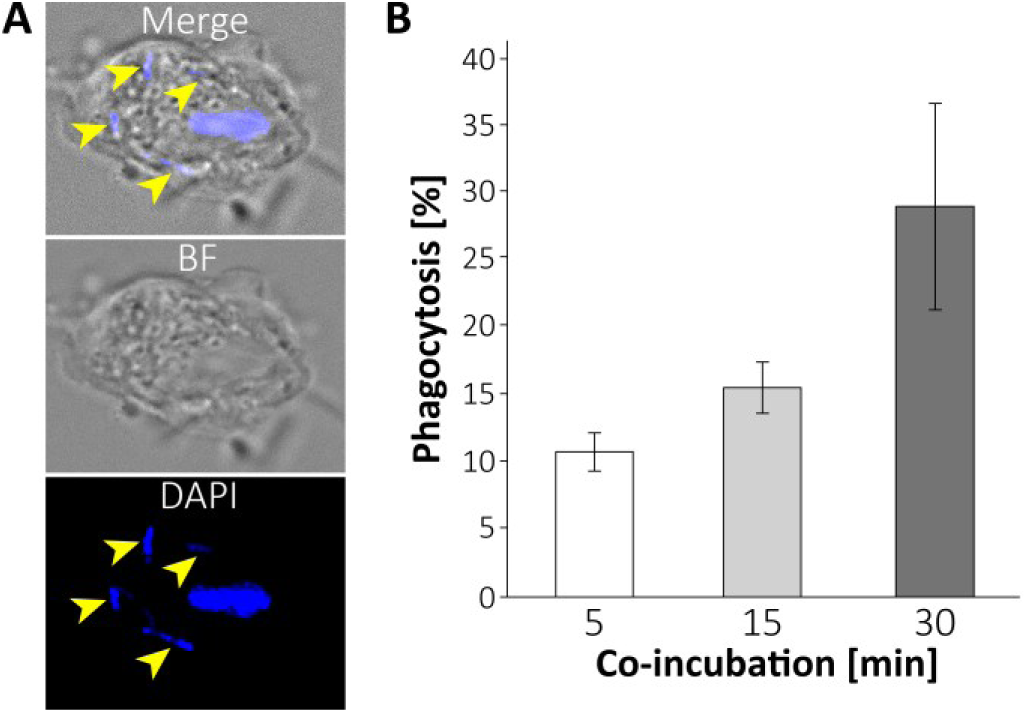
Phagocytic activity of *T. vaginalis* (TV) to *Lactobacillus jensenii* (LJ). TV was co-incubated with LJ at 37°C for 5, 15, and 30 min. **A.** Representative images of a TV cell showing phagocytosed LJ after 15 min of co-incubation. The arrows indicate internalized lactobacilli. BF, Brightfield. **B.** Phagocytosis over time. After 30 min of co-incubation, about 30% of TV cells had ingested LJ. Error bars represent standard deviation. N = 10^2^ cells/replicate, three replicates/sample.

**Figure 2.**
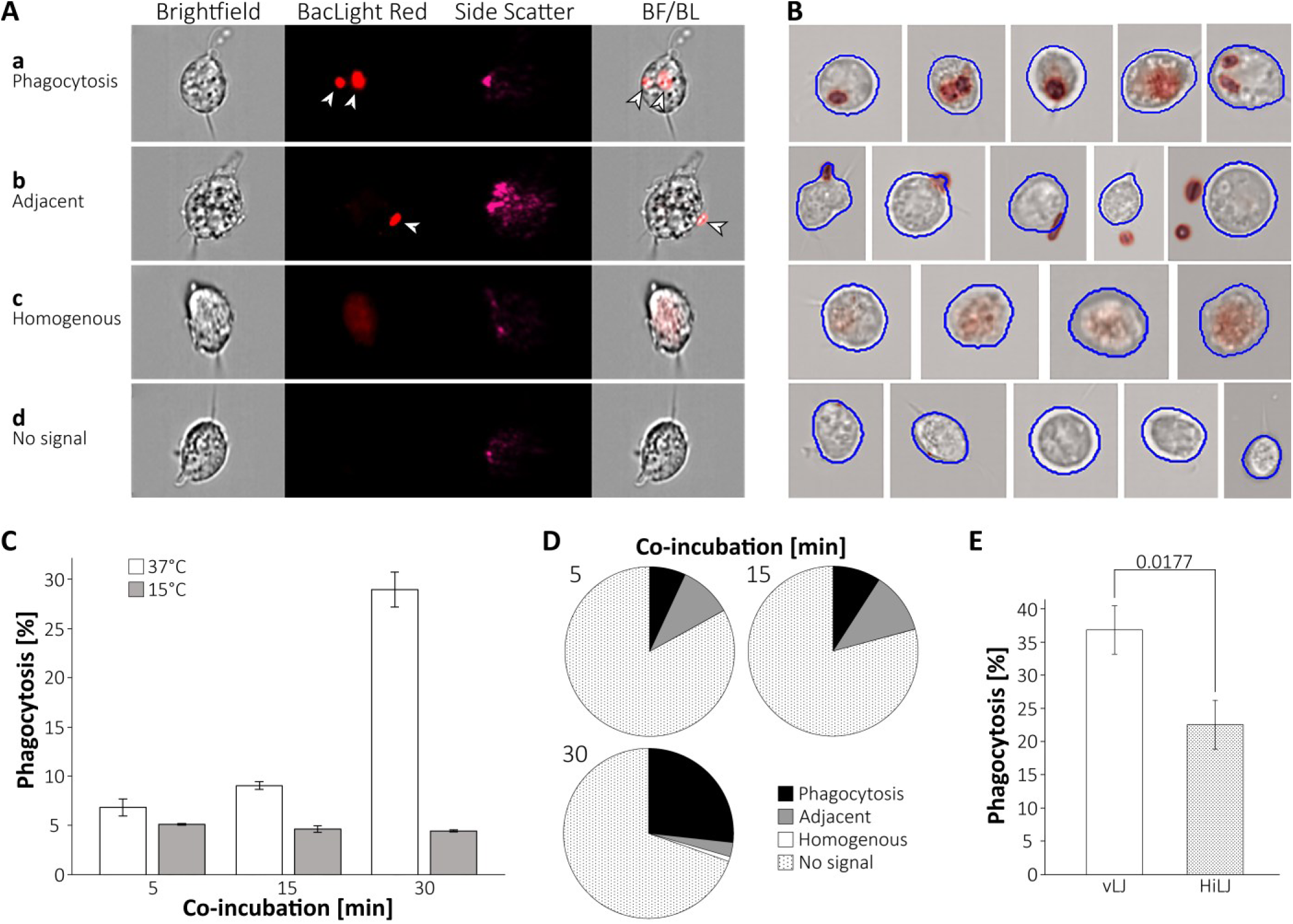
Imaging flow cytometry analysis of *T. vaginalis* (TV) phagocytosis confirmed high phagocytic activity to *L. jensenii* (LJ). TV was co-incubated with BacLight Red-labelled LJ at 37°C for 5, 15, and 30 min. **A.** The analysis revealed four groups based on the BacLight Red signal: a. Phagocytosis, with internalized LJ signal; b. Adjacent, with LJ signal on or close to TV’s cell surface; c. Homogenous, with BacLight Red signal homogenously distributed across the TV cell; d. No BacLight Red signal. LJ cells are indicated with an arrow. BF, Brightfield; BL, BacLight Red. **B.** Automated segmentation of images acquired through imaging flow cytometry. Rows indicate the groups as shown in A. **C.** Co-incubation of TV with LJ showed time-dependent increasing phagocytosis, with an average of 30% after 30 min at 37 °C, whereas phagocytic activity remained low in the control at 15 °C (5%). Error bars represent standard deviation. N = 1.2 x 10^3^ cells/replicate, three replicates/sample. A two-way ANOVA followed by Tukey’s multiple comparison test was performed and revealed a significant result (p < 0.001). **D.** Dynamics of groups at 37°C over time. While the phagocytosis group grew significantly over time, TV cells with homogenous BacLight Red signal stayed below 2%. **E.** Co-incubation with either viable LJ (vLJ) or heat-inactivated LJ (HiLJ) for 30 min revealed that TV preferentially phagocytoses vLJ. Error bars represent standard deviation. The number above the bars indicates *P*-value for the unpaired t-test. N = 2.5 x 10^3^ cells/replicate, three replicates/sample.

### *T. vaginalis* ingests *L. jensenii* via a pseudopodia-independent process

To visually investigate the LJ phagocytosis by TV, microscopy images were taken following TV-LJ co-incubation (Fig. 3). In fluorescence microscopy, LJ appeared as long rods that co-localized with the endolysosomal marker Rab7 (29), demonstrating the engulfment into TV phagolysosomal compartment (Fig. 3A). Typically, multiple lactobacilli were observed in an individual TV cell. In time-dependent experiments, LJ phagocytosis was observed using transmission electron microscopy (TEM). Lactobacilli appeared inside TV cells as early as five minutes of co-incubation (Fig. 3B). Up to four LJ cells were observed in a single TV cell, each of them localizing to individual phagolysosomes. Interestingly, the formation of pseudopodia during phagocytosis was not observed. Thus, to investigate the cell surface structures during phagocytosis, scanning electron microscopy (SEM) was employed (Fig. 3C). This analysis revealed that LJ is not phagocytosed by classical cell membrane protrusions. Instead, LJ initially attached to the TV cell via its tip and subsequently was pulled into the TV cells through a narrow opening reminiscent of a sinking type of phagocytosis (30–32). In contrast, when TV was co-incubated with *Candida albicans*, no phagocytosis was observed after 15 and 60 min of co-incubation. However, after 120 min of co-incubation, TV formed classical pseudopodia to engulf the cell (Fig. 3C).

**Figure 3.**
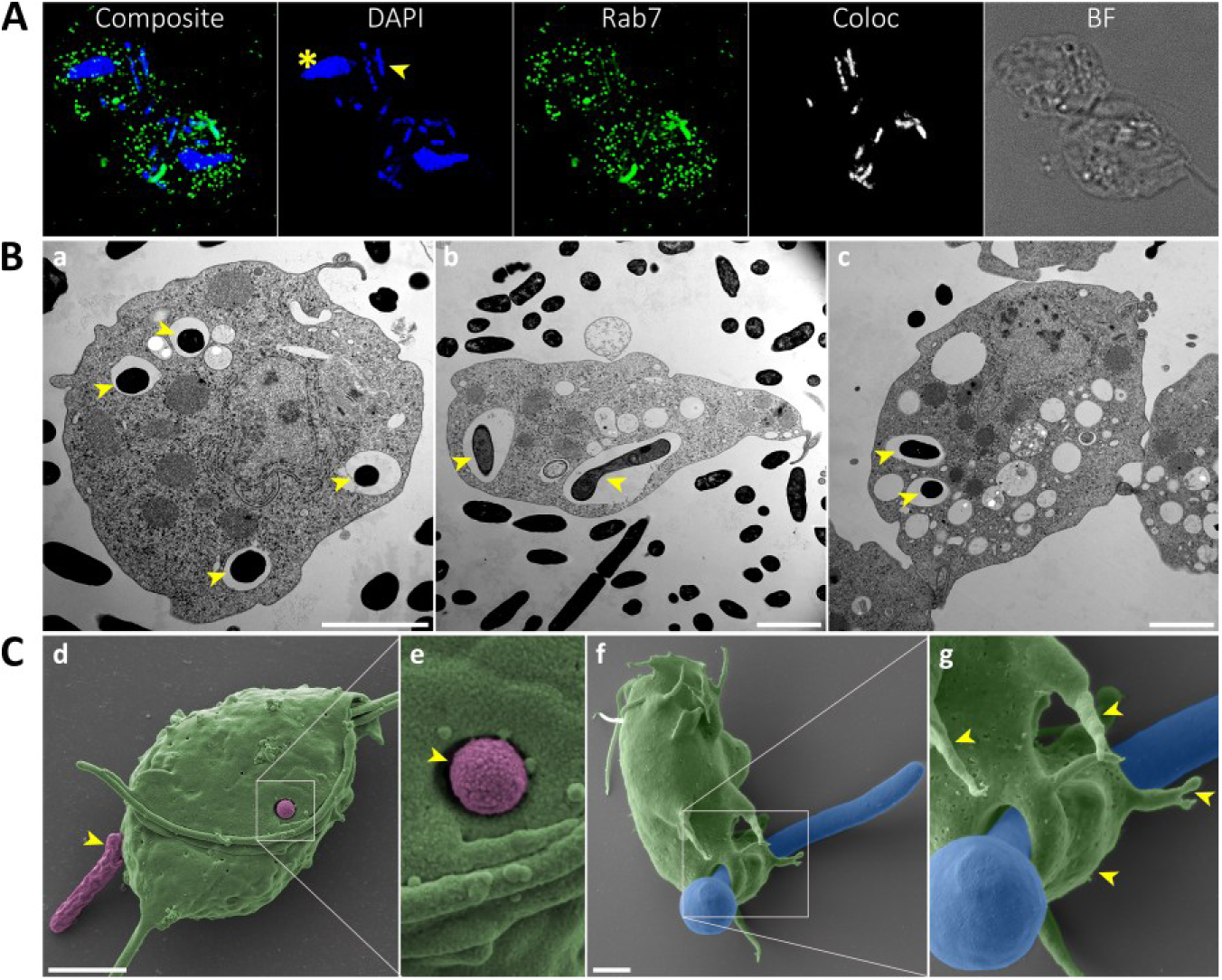
Rapid engulfment of *L. jensenii* (LJ) by *T. vaginalis* (TV) via pseudopodia-independent phagocytosis. **A.** Fluorescence microscopy was performed after 30 min of co-incubation of TV with LJ. Rab7 (green) was used as an endo-lysosomal marker and stained with mouse monoclonal anti(α)-HA antibody and Alexa Fluor 488 donkey α-mouse. DAPI (blue) was used to visualize both lactobacilli and *Trichomonas* nuclei. The arrow points at an exemplary LJ cell, whereas the asterisk indicates a TV nucleus. The colocalization of lactobacilli with Rab7 is shown in the colocalization channel (Coloc). BF, Brightfield. **B.** Transmission electron microscopy (a-c). Phagocytosed lactobacilli (black) were observed in individual phagolysosomes (arrows) after 5 min of co-incubation. Scale bar = 2 µm. **C.** Scanning electron microscopy revealed pseudopodia-independent engulfment of LJ (pink) by TV cells (green). (d) The arrow indicates attachment of LJ to the TV surface. (e) The arrow points at LJ engulfment via a narrow opening. (f-g) Phagocytosis of *Candida albicans* (blue) by TV (green) was observed after 120 min of co-incubation and served as a control for pseudopodia-dependent phagocytosis (arrows). Scale bar = 2 µm.

### Presence of *L. jensenii* enhances *T. vaginalis* secretion of a specific set of proteins

Next, we analyzed the secretome upon TV-LJ co-incubation for 10, 30, and 60 min at 37°C and on ice as a negative control. The induced secretome was compared with the constitutive secretome in the absence of LJ. MS analysis yielded 933 TV proteins in the co-incubation samples (Dataset S1) and 954 identifications in the control without lactobacilli (Dataset S2). Of these, 85 proteins revealed a time-dependent increase in conditioned medium and met the criteria defined for secreted proteins (Fig. S1). The increased secretion stimulated by LJ was found for 27 proteins (TvIS). The significance was tested based on differences in the LFQ values (Kruskal-Wallis test, p < 0.05) and secretion scores (t-test, p < 0.05) (Fig. S2 and Fig. S3, respectively).

The TvIS proteins showed differences in secretion dynamics, based on which three distinct secretion groups were assigned (Fig. 4). Proteins in group 2 exhibited the strongest time-dependent upregulation in secretion, indicated by the highest z-score increase. This group comprises three degradative enzymes: a leishmanolysin-like metallopeptidase (GP63), which is a surface-anchored protease, a putative trehalase family protein, and glycosyl hydrolase 25 lytc-like domain-containing protein (TvGH25). Furthermore, it includes six paralogs of *Trichomonas* beta-sandwich repeat (TBSR) proteins. TBSRs are proteins that resemble metazoan cadherins, which are cell surface proteins involved in cell-cell interaction (10, 33, 34). They are comprised of a single transmembrane domain close to the C-terminus, a glycosyl hydrolase-like domain, and beta-sandwich repeat structures (Fig. S4). Another TBSR protein [TBSR1, previously described as BAP1 (35)] clustered in group 1. This group also includes an α-1,6-glucosidase and an α-amylase, both of which belong to the glycogen-degrading class of enzymes. Other members of this group are TvSaplip1, a protein of the pore-forming toxin family (36–38), and a DNase I-like protein. Group 3 includes two peptidases (Cathepsin D-like and Papain-like peptidase), a polymorphic outer membrane protein (PMP), as well as proteins related to the nucleus and hydrogenosomes (Fig. 4).

**Figure 4.**
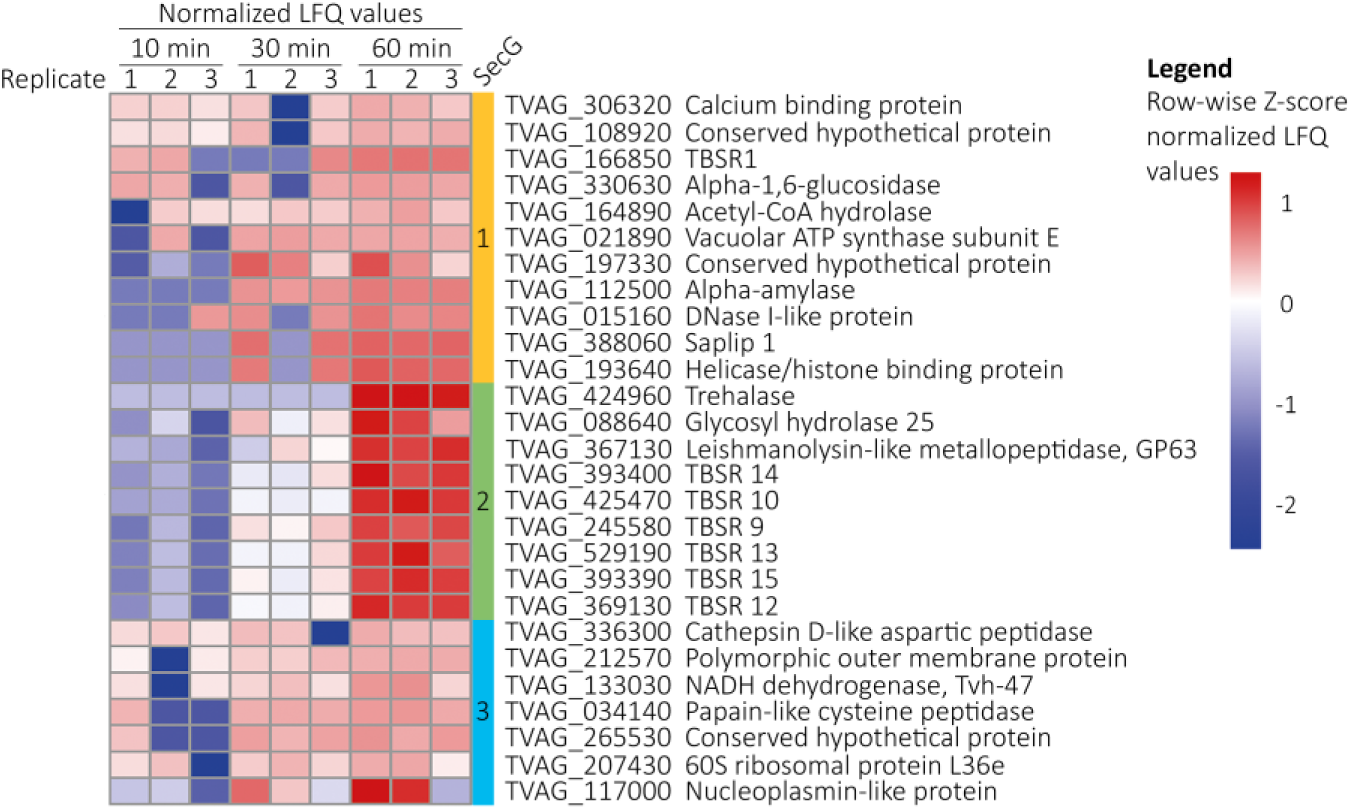
*T. vaginalis* (TV) enhances the secretion of a specific protein subset in response to *L. jensenii* (LJ). The heatmap showed TV proteins with a significantly increased secretion upon co-incubation of TV with LJ. Each row represents the relative amount of proteins [z-score normalized label-free quantification (LFQ) values] determined by mass spectrometry in three biological replicates for each time point. The proteins were clustered into three secretion groups (SecG) based on the Pearson correlation of normalized LFQ values. The subset of proteins was filtered based on the statistical evaluation given in Figures S1, S2, and S3.

### Real-time quantitative PCR reveals upregulation of TvIS proteins on RNA level

To validate the proteomic results, we selected three proteins to analyze their gene expression by Real-time quantitative PCR (RT-qPCR) in response to co-incubation with LJ: TBSR1-Bap1, with a rather constant time-dependent increase on the proteomic level (Group 1), and two proteins of Group 2, TBSR12, and TvGH25, with marked upregulation after 60 min (Fig. 4). All three genes showed upregulation over time relative to the control without LJ (Fig. 5). TBSR1 expression increased moderately at an approximately constant rate as was observed on the proteomic level. TBSR12 expression initially did not change significantly, but increased by 50% after 60 min of co-incubation. Similarly, significant upregulation after 60 min of co-incubation was observed for TvGH25. The dynamics of RNA level upregulation for TBSR1, TBSR12, and TvGH25 thus correspond to the proteomic data and further support their active role in TV response to LJ presence.

**Figure 5.**
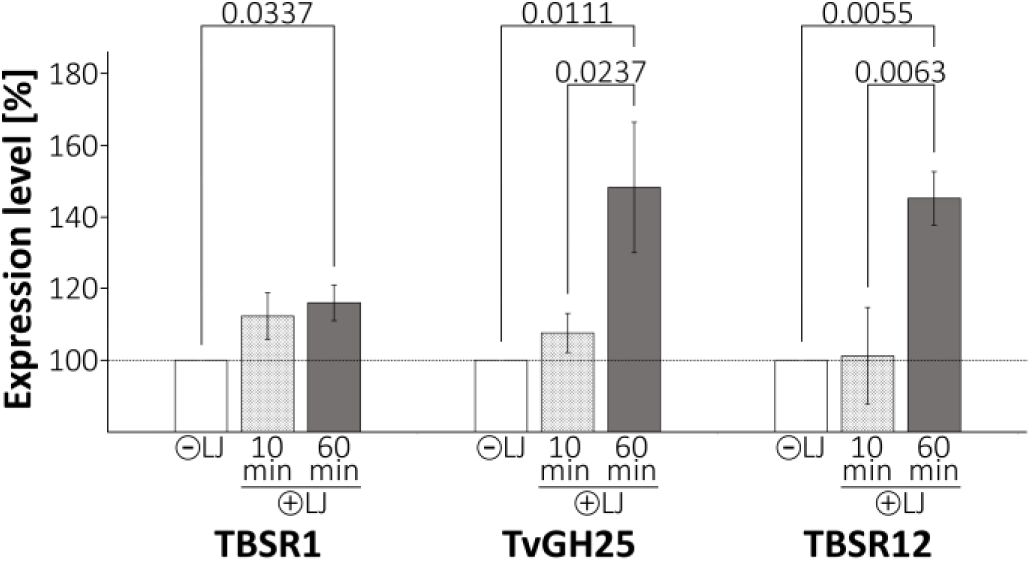
Increased RNA levels of TBSR1, TBSR12, and TvGH25 correspond to their increased secretion in response to *T. vaginalis* -*L. jensenii* (TV-LJ) co-incubation. Relative RNA levels in TV were determined by qRT-PCR for selected genes after co-incubation of TV with LJ for 0, 10, and 60 min. Error bars represent standard deviation. N = 3 replicates/sample. Numbers above bars indicate *P*-values for one-way ANOVA followed by Tukey’s multiple comparison test. TvGH25, Glycosyl hydrolase 25; TBSR, *Trichomonas* beta-sandwich repeat-protein.

### TvGH25 was acquired by lateral gene transfer

The glycosyl hydrolase family 25 (GH25) represents a prokaryotic type of lysozyme, although its members have also been found in viruses and some eukaryotes (mainly fungi) (39). These enzymes cleave the bacterial cell envelope polymer, peptidoglycan; thus, the identification of putative GH25 in TV among TvIS proteins designates it as an excellent candidate for antibacterial activity. Searches in the TV genome revealed four TvGH25 paralogs (Fig. S5), and multiple GH25 genes were found in the genomes of all members of the Trichomonadidae group of Parabasalia, but not in *Tritrichomonas foetus* (Table S1). The Trichomonadidae sequences were interleaved within a prokaryotic branch of mostly firmicutes and a bacteriophage (Fig. 6). Based on this topology, it could be inferred that the Trichomonadidae ancestor acquired GH25 most likely via lateral gene transfer (LGT) from prokaryotes. The analysis also suggests an independent acquisition of GH25 by *Dientamoeba fragilis*, another member of Parabasalia. The second major clade included sequences found in mostly unicellular eukaryotes of Metamonada, Discoba, Amoebozoa, Stramenopiles, and fungi, as well as members of invertebrates (Nematoda, Rotifera, and sponges). Importantly, the common feature of these organisms is that they feed on bacteria (38).

**Figure 6.**
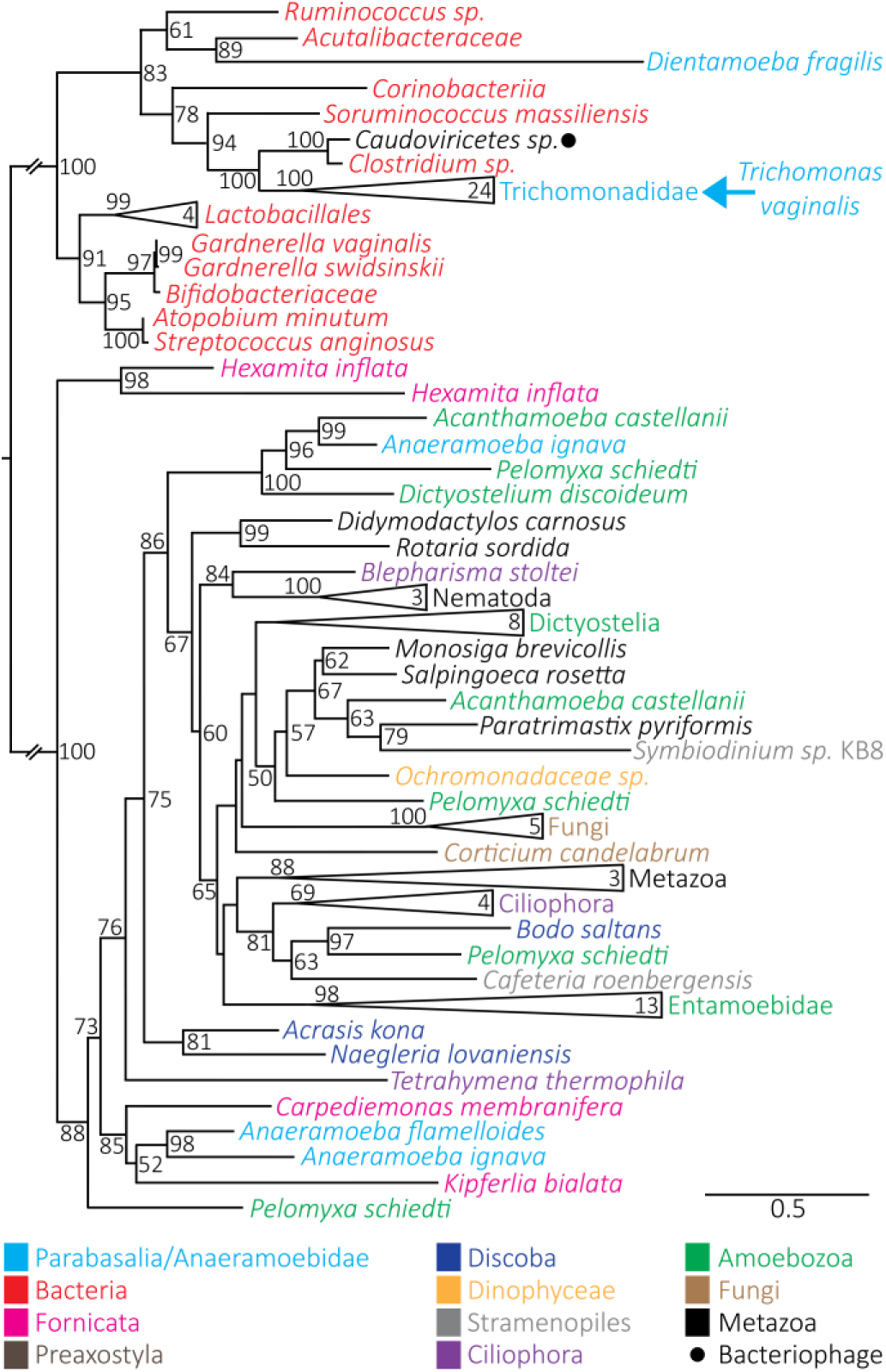
Phylogenetic analysis indicates lateral gene transfer of TvGH25 from bacteria to the Trichomonadidae ancestor. Taxa are colour-coded. Nonparametric bootstrap values are indicated at nodes; values below 50 are omitted. Numbers of sequences in collapsed branches are indicated within triangles. The analyzed sequences are listed in Table S1.

### TvGH25 displays muramidase activity

GH25 antibacterial activity is based on the cleavage of the β-1,4-glycosidic bond between N-acetylmuramic acid (Mur*N*Ac) and N-acetylglucosamine (Glc*N*Ac) in bacterial peptidoglycan (PG). To determine the activity of TvGH25, we isolated PG from *L. gasseri* ATCC 9857 and *E. coli* BW25113Δ6LDT (40). These substrates were treated with recombinant TvGH25 expressed in *E. coli* and the bacterial muramidase cellosyl from *Streptomyces ceolicolor* (41). HPLC analysis of the digested substrates revealed an identical pattern of muropeptides for TvGH25 and cellosyl (Fig. 7A and B). In contrast, TvGH25 with mutated catalytic active site Asp100Ala and Glu102Gln did not release any muropeptides (Fig. 7A). These results demonstrated the muramidase activity of TvGH25 against bacterial peptidoglycan, a major cell wall component.

**Figure 7.**
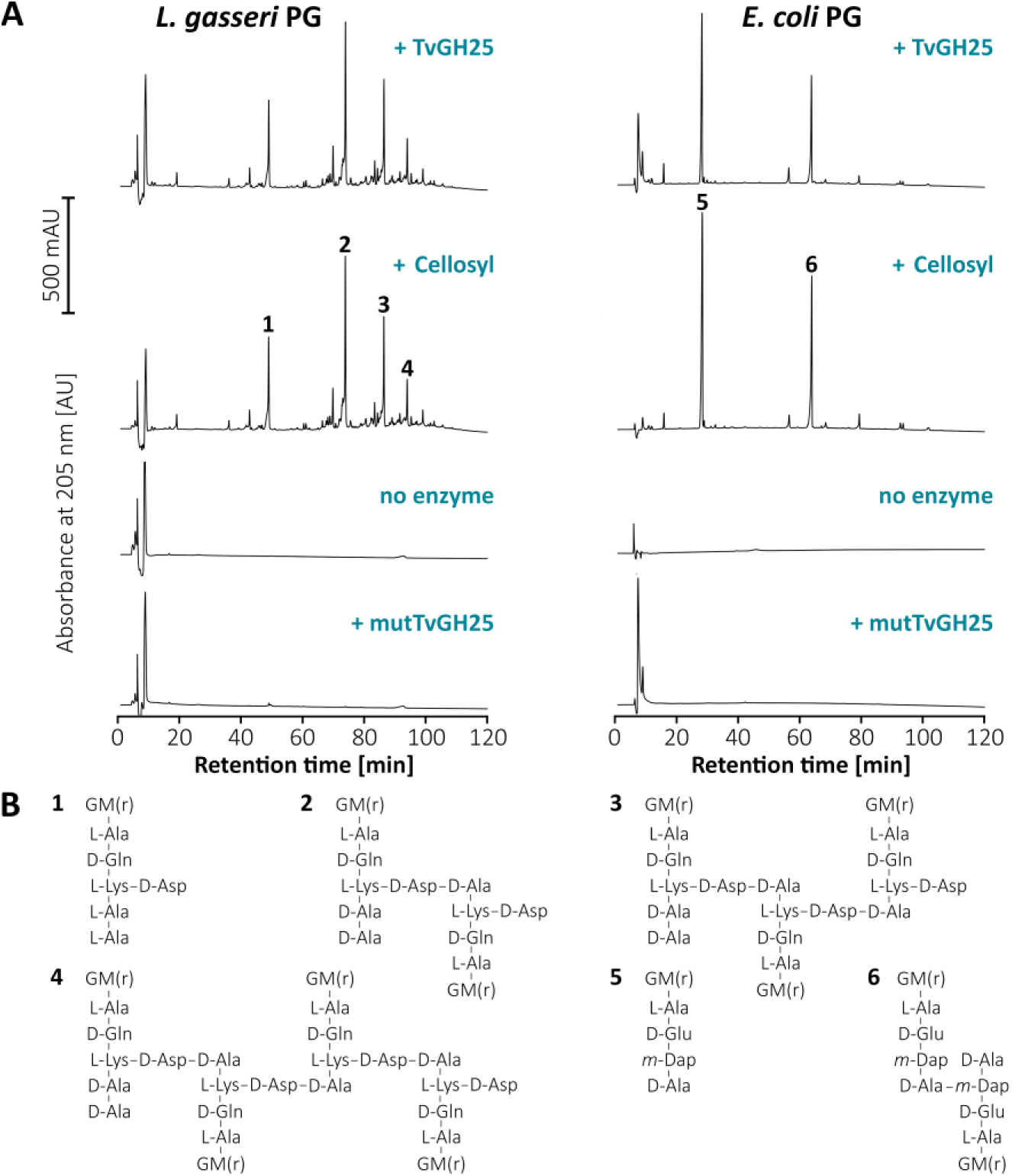
Muramidase activity of TvGH25. **A.** Peptidoglycan (PG) from *L. gasseri* ATCC 9857 or *E. coli* BW25113Δ6LDT (homogeneous tetrapeptides-rich) was incubated with recombinant TvGH25, mutTvGH25, the muramidase cellosyl, or without enzyme, followed by reduction with sodium borohydride and separation of muropeptides by HPLC. Cellosyl and TvGH25 released muropeptides, while no muropeptides were released by mutTvGH25 or when the enzyme was omitted in the reaction. Numbers above peaks indicate main muropeptides released from the PG, detailed in (B). **B.** Structures of the main muropeptides released from the PG. M(r), N-acetylmuramitol; G, N-acetylglucosamine; L-Ala, L-alanine; D-Ala, D-alanine; D-Glu, D-glutamic acid; D-Gln, D-glutamine; L-Lys, L-lysine, D-Asp, D-aspartic acid; m-Dap, meso-diaminopimelic acid.

### Impact of TvGH25 on *L. jensenii*

To evaluate the antimicrobial activity of TvGH25 against LJ, we overexpressed TvGH25 in TV under the control of a strong promoter (42). Immunofluorescence microscopy confirmed that recombinant TvGH25 localized to the endoplasmic reticulum, Golgi apparatus associated with the parabasal filament, and multiple vesicles, consistent with the classical ER-Golgi secretory pathway (Fig. 8A). Western blot analysis of TV conditioned medium further verified secretion of recombinant TvGH25 (Fig. 8B).

**Figure 8.**
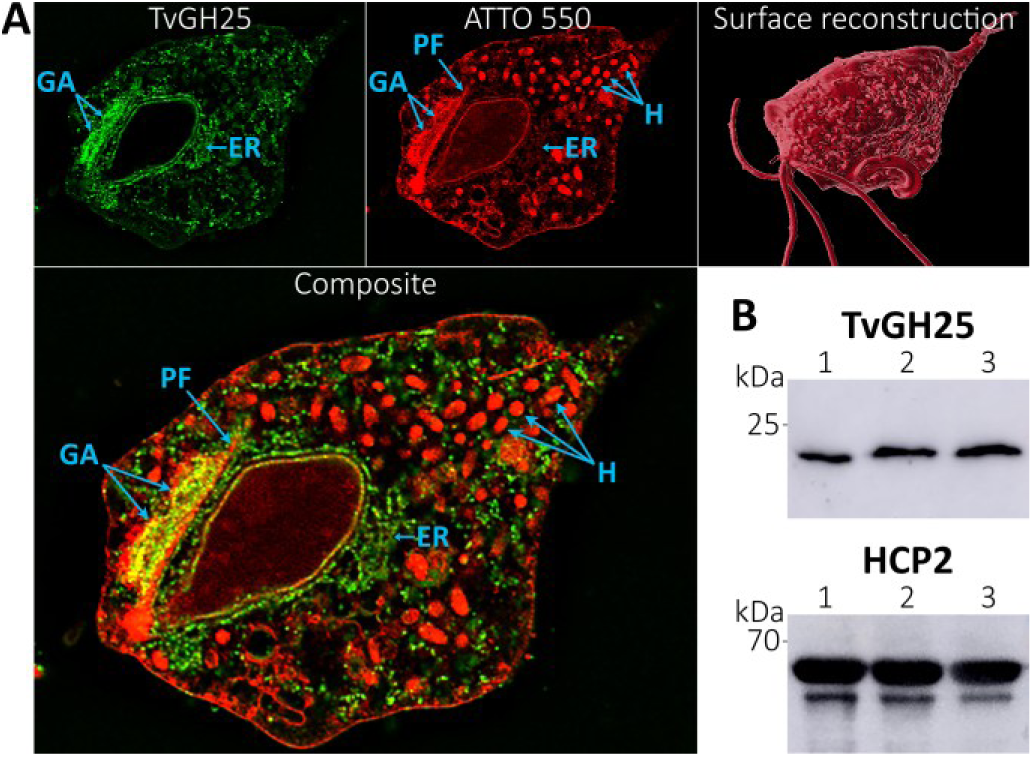
ER-Golgi secretion of recombinant TvGH25. **A.** Intracellular localization of V5-tagged TvGH25 by expansion microscopy. TvGH25 (green) was visualized using mouse monoclonal anti(α)-V5 antibody and donkey α-mouse antibody conjugated to Alexa Fluor 488. The fluorescent probe ATTO 550 NHS ester was used to label protein-rich cell structures, visualizing vesicles, endoplasmic reticulum (ER), the Golgi apparatus (GA), and hydrogenosomes (H). TvGH25 strongly co-localized with the fluorescent probe in the Golgi apparatus (arrow), supported by the parabasal filament (PF), as well as in the ER, and multiple tiny vesicles scattered within the cytosol. **B.** Western Blot analysis of TV conditioned medium showing secreted recombinant TvGH25 after 10 min of incubation. Secreted TvGH25 was detected using α-V5 antibody. Hydrogenosomal HCP2 was used as a loading control. Numbers in lanes indicate biological replicates. TvGH25, Glycosyl hydrolase 25; BF, Brightfield; HCP2, Hybrid Cluster Protein 2.

Next, LJ cells were co-incubated with either TV overexpressing a recombinant TvGH25 or wild-type (WT) parasites, and colony-forming units (CFUs) were compared to the untreated LJ control cultured without trichomonads. Aliquots of the co-incubations were plated on agar after 10 to 300 minutes (fig. 9A), and CFUs were counted following overnight incubation at 37°C. In the co-incubations with both WT and TvGH25-overexpressing TV, significantly lower CFU levels were observed from 180 minutes (Fig. 9B). After 300 minutes, CFU levels of the co-incubation with WT TV were at approximately 70% compared to the control, whereas CFU levels of the co-incubation with the TvGH25-overexpressing strain reached only 30% of control levels (Fig. 9C). To confirm the specific contribution of TvGH25, LJ was co-incubated with a control TV strain expressing the recombinant functionally unrelated protein HCP1, which did not affect LJ CFUs (Fig. 9B,C). These results demonstrate that TV reduces LJ growth/viability, and that this effect is enhanced by overexpression of secreted TvGH25, supporting its role in its antimicrobial activity against LJ.

**Figure 9.**
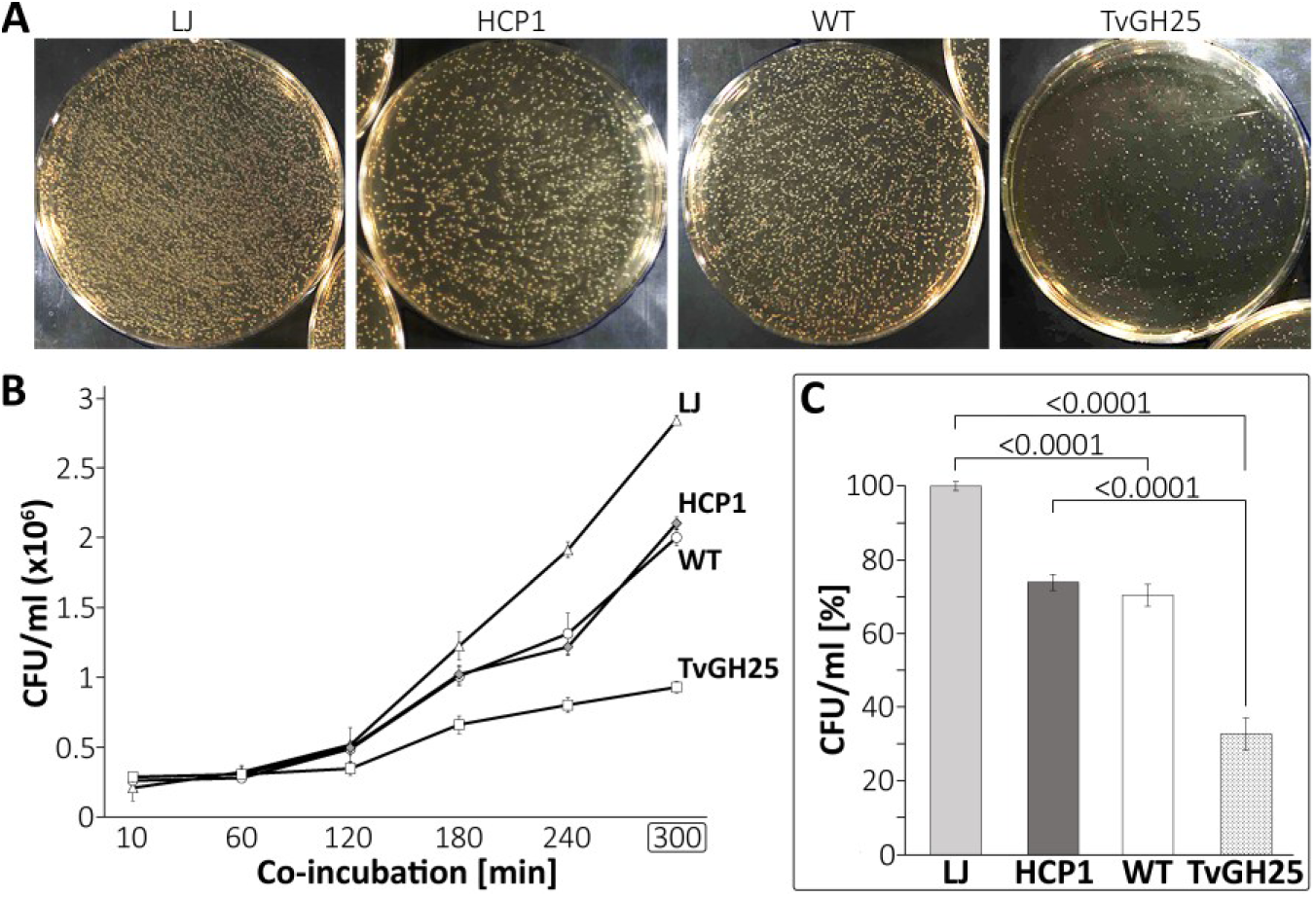
TvGH25 overexpression impacts *L. jensenii* (LJ) population. LJ cells were co-incubated for up to 300 min with either wild-type *T. vaginalis* (WT TV) or TV overexpressing V5-tagged TvGH25. As controls, lactobacilli were incubated axenically (LJ) or co-incubated with TV overexpressing hybrid cluster protein 1 (HCP1). Subsequently, diluted aliquots of lactobacilli were seeded on plates and grown overnight. **A.** Representative images of plates after 300 min of co-incubation are shown. Significant differences in bacterial growth are visible in the co-incubations compared with the axenic incubation (LJ). Aliquots were diluted 1:10^2^ prior to plating. **B.** Counts of LJ colony forming units (CFUs) showed that, after 180 minutes of co-incubation, significant differences began to emerge between the co-incubations. Specifically, the CFUs of LJ co-incubated with TV overexpressing TvGH25 remained significantly lower compared to the co-incubation with WT or in the axenic incubation of LJ. Aliquots were diluted 1:10^3^ prior to plating for accurate counts. Error bars reflect standard deviation. N = 3 replicates/sample. **C.** Differences in CFUs after 300 min of co-incubation normalized to axenic LJ. CFU counts after co-incubation with TvGH25 overexpressing TV were only 33% of axenic LJ. Error bars reflect standard deviation. Numbers above bars indicate *P*-values for one-way ANOVA followed by Tukey’s multiple comparison test.

## Discussion

Parasitic protists, such as TV, secrete proteins that modulate their environment and interact with host cells to facilitate survival and establish infection. However, little is known about how the protein secretion by parasitic protists is influenced by bacterial members of the host microbiota. In our study, we employed a simplified *in vitro* system comprising TV and LJ, a dominant protective *Lactobacillus* species associated with CST-V, to investigate the impact of LJ on the complex secretome profile of TV. Our data demonstrated a rapid and robust response of TV to the presence of LJ by an increased secretion of 27 TvIS proteins that have the potential to interact with and damage bacteria. Among these TvIS proteins is TvGH25, which belongs to a large family of diverse enzymes called lysozymes, all of which exhibit muramidase activity. In metazoans, including the human host of TV, secretion of lysozyme represents a key defense mechanism for killing bacteria (43). However, TvGH25 is a typical bacterial enzyme that is structurally unrelated to the human lysozyme. Indeed, our phylogenetic analysis indicates that TvGH25 was acquired by LGT from bacteria, which extends the previous report suggesting broad intradomain or interdomain transfer of GH25 genes (39). The ancestral role of GH25 in bacteria was likely the remodeling of the cell wall during cell reproduction (44), while GH25 that was transferred outside of bacteria was repurposed for antibacterial action (39). Our data indicate that this is also the case for TvGH25. We demonstrated TvGH25 activity against purified peptidoglycans and the production of defined muropeptide fragments, as those generated by cellosyl. Notably, TvGH25 overexpression in TV led to significantly reduced LJ viability, providing functional evidence that TvGH25 contributes to the antibacterial activity. A key role of eukaryotic GH25 in bacterial killing and digestion is further supported by the distribution of GH25 in free-living as well as parasitic protists. Our homology searches across eukaryotic lineages identified this enzyme exclusively in organisms that prey on bacteria. For example, in Parabasalia, GH25 is present in TV and related species of the Trichomonadidae group, which are all bacteria-phagocytosing organisms. In contrast, *Tritrichomonas foetus* of Tritrichomonadidae, which acquires nutrients dominantly via fluid-phase and receptor-mediated endocytosis with a lower capacity to ingest particles (45), lacks this enzyme. Similarly, GH25 was acquired by the diplomonad *Hexamita*, which feeds on bacteria using well-developed cytostomes, while it is absent in closely related *Giardia*, which lack cytostomes and acquire nutrients via a specific peripheral endocytic compartment (46–48). Among metazoans, GH25 is present, for example, in sponges that feed on bacteria by water filtering through their bodies. Altogether, increased secretion of TvGH25 in the presence of LJ and its muramidase activity indicates that this enzyme belongs to the key components of TV antibacterial tools.

Previously, NlpC/P60-type peptidoglycan hydrolases were shown to be in the arsenal of TV antibacterial weapons (11, 49). Like TvGH25, this enzyme targets bacterial peptidoglycan; as a cysteine peptidase, it cleaves a peptide bond within the stem peptides of peptidoglycan (11, 49). Thus, secretion of both NlpC/P60 and TvGH25 is expected to have a synergistic effect to destroy the bacterial cell wall. In line with this view, the transcription of two NlpC/P60 paralogs is upregulated in the presence of *E. coli*, and TV cells expressing a recombinant NlpC/P60 were more efficient in reducing bacterial counts (11). The TV genome contains nine NlpC/P60 genes of two classes (A and B) (11). Intriguingly, only a single NlpC/P60 paralog (NlpC/P60-A2) has been found in the constitutive TV secretome (10), while none of them was found among TvIS in this study. This result is unexpected and most likely reflects the selectivity of TV response to different bacteria. While in our experiment TV was challenged with Gram-positive LJ, increased NlpC/P60 transcription was observed upon challenge with Gram-negative *E. coli* (11). In addition, the result of TV-bacteria interaction could also be strain-specific (50, 51). Thus, the interplay between TvGH25 and the NlpC/P60 enzyme needs further study to elucidate their specific role in TV-bacteria interactions. Of note, similar to TvGH25, NlpC/P60A and NlpC/P60B were also acquired by LGT from bacteria (49), highlighting how the coevolution of *Trichomonas* ancestors and specific microbiota shaped the genome of these parasites, providing selective advantages for inhabiting their niche.

TvIS proteins included three proteases: aspartic protease Cathepsin D (CatD), metalloprotease (M8 family, GP63), and papain-like cysteine peptidase. In addition, at least 13 proteases of various catalytic classes, such as M8, M24, and M60 metalloproteases, serine and papain-like cysteine peptidases, are constitutively secreted (10). Although involvement of proteases in TV virulence is well established (52–54), their role in TV-bacteria interaction is not known. The surface of lactobacilli, including LJ, is covered by S-layer proteins and aggregation-promoting factors (APFs) that are noncovalently attached to peptidoglycan and form a protective barrier (18). Thus, proteolytic cleavage of this barrier likely precedes the digestion of peptidoglycan. The observed increase in the secretion of three proteases upon LJ co-incubation makes them interesting candidates for such a function.

Another TvIS protein is TvSaplip-1, a member of the saposin-like protein (Saplip) family. Saplip proteins, such as Ameobapores and Naegleriapores, studied in *Entamoeba histolytica* and *Naegleria fowleri*, respectively, can lyse both bacterial and eukaryotic cells by forming pores in their membranes (38, 55). TV possesses twelve Saplip paralogs, with the function of TvSaplip12 having been studied previously, where its antibacterial activity was demonstrated (56). Interestingly, unlike TvSaplip12, expression of TvSaplip-1 was stimulated upon contact with LJ, but not with human red blood cells (56). Such a selective response further supports a possible role of TvSaplip-1 in TV-LJ interaction.

Besides proteases, the second largest group of TvIS proteins is TBSR membrane cadherin-like proteins. TBSRs have previously been found in the surface proteome (57), lysosomal proteome (29), and the constitutive secretome (10) of TV (Table 1). Such a distribution suggests that TBSRs are present on the outer TV membrane and could be either engulfed into lysosomes or released to the environment. Indeed, one of these, TVAG_393390 [named CLP, (58)], has been shown to localize to the TV membrane and possesses a putative cleavage site for membrane rhomboid protease 1 (TvROM1) to be cleaved off from the membrane (58, 59). Furthermore, the overexpression of CLP increased the adherence of TV to the host cells and wild-type trichomonads (58). The strong upregulation of multiple TBSRs in response to LJ presence suggests that TBSRs are involved in adhesive functions, also in parasite-microbe interactions, and possibly contribute to phagocytosis of lactobacilli. Further investigations on this class of proteins are necessary to identify their ligands on the *Lactobacillus* surface and elucidate their specific role in TV-microbe interactions.

**Table 1.**
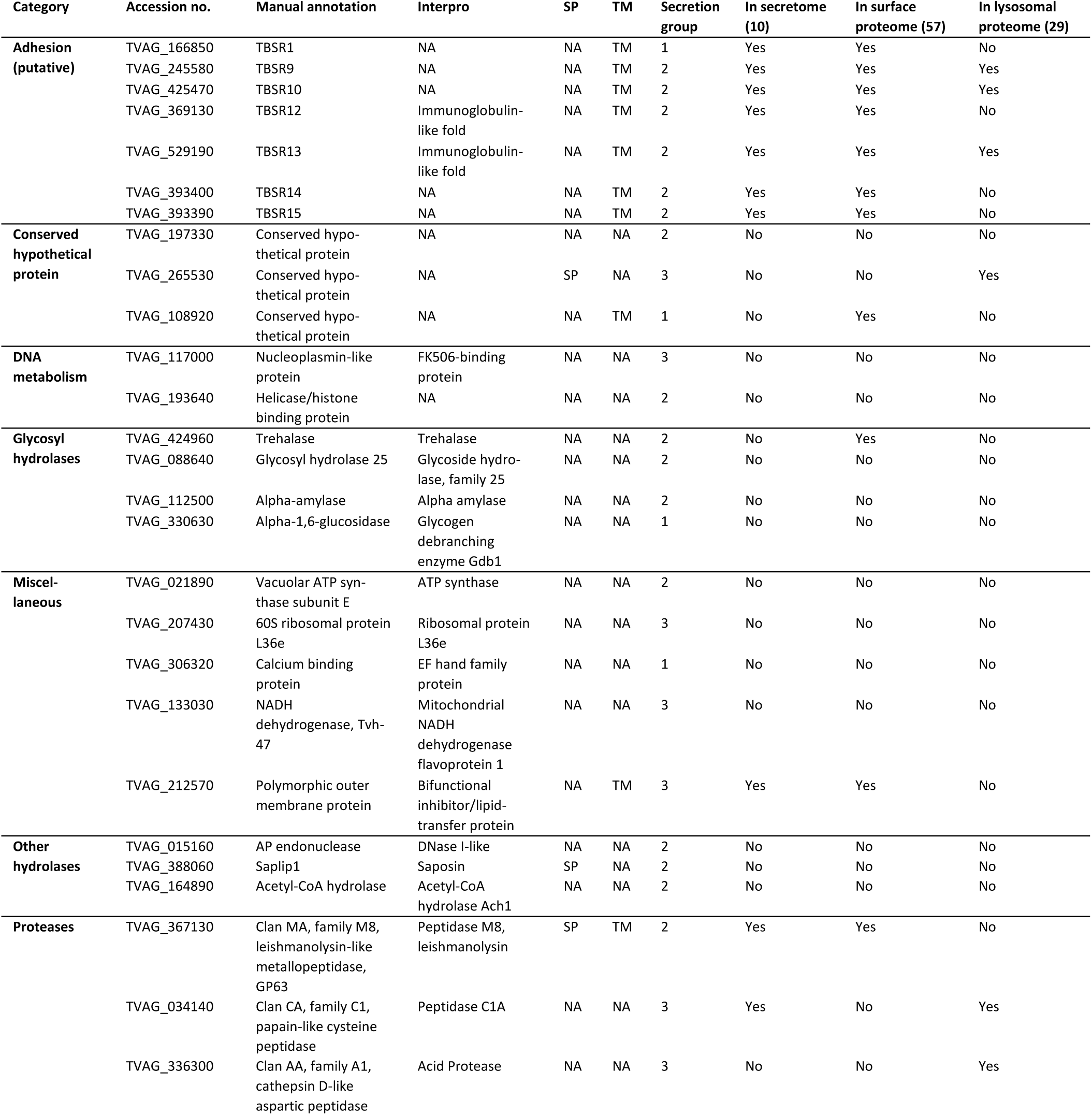
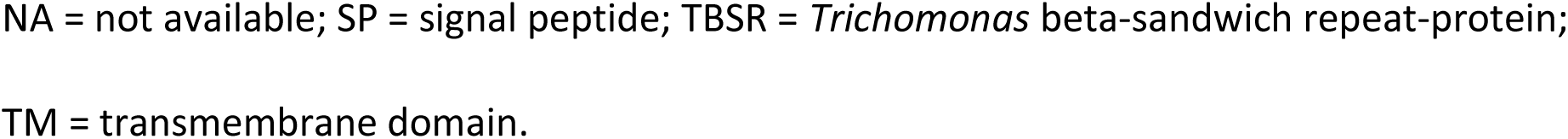
Subset of proteins with significantly increased secretion by *T. vaginalis* in response to co-incubation with *L. jensenii*. The proteins were categorized based on their assigned function.

| Category | Accession no. | Manual annotation | Interpro | SP | TM | Secretion group | In secretome (10) | In surface proteome (57) | In lysosomal proteome (29) |
| --- | --- | --- | --- | --- | --- | --- | --- | --- | --- |
| **Adhesion (putative)** | TVAG_166850 | TBSR1 | NA | NA | TM | 1 | Yes | Yes | No |
|  | TVAG_245580 | TBSR9 | NA | NA | TM | 2 | Yes | Yes | Yes |
|  | TVAG_425470 | TBSR10 | NA | NA | TM | 2 | Yes | Yes | Yes |
|  | TVAG_369130 | TBSR12 | Immunoglobulin-like fold | NA | TM | 2 | Yes | Yes | No |
|  | TVAG_529190 | TBSR13 | Immunoglobulin-like fold | NA | TM | 2 | Yes | Yes | Yes |
|  | TVAG_393400 | TBSR14 | NA | NA | TM | 2 | Yes | Yes | No |
|  | TVAG_393390 | TBSR15 | NA | NA | TM | 2 | Yes | Yes | No |
| **Conserved hypothetical protein** | TVAG_197330 | Conserved hypothetical protein | NA | NA | NA | 2 | No | No | No |
|  | TVAG_265530 | Conserved hypothetical protein | NA | SP | NA | 3 | No | No | Yes |
|  | TVAG_108920 | Conserved hypothetical protein | NA | NA | TM | 1 | No | Yes | No |
| **DNA metabolism** | TVAG_117000 | Nucleoplasmin-like protein | FK506-binding protein | NA | NA | 3 | No | No | No |
|  | TVAG_193640 | Helicase/histone binding protein | NA | NA | NA | 2 | No | No | No |
| **Glycosyl hydrolases** | TVAG_424960 | Trehalase | Trehalase | NA | NA | 2 | No | Yes | No |
|  | TVAG_088640 | Glycosyl hydrolase 25 | Glycoside hydrolase, family 25 | NA | NA | 2 | No | No | No |
|  | TVAG_112500 | Alpha-amylase | Alpha amylase | NA | NA | 2 | No | No | No |
|  | TVAG_330630 | Alpha-1,6-glucosidase | Glycogen debranching enzyme Gdb1 | NA | NA | 1 | No | No | No |
| **Miscellaneous** | TVAG_021890 | Vacuolar ATP synthase subunit E | ATP synthase | NA | NA | 2 | No | No | No |
|  | TVAG_207430 | 60S ribosomal protein L36e | Ribosomal protein L36e | NA | NA | 3 | No | No | No |
|  | TVAG_306320 | Calcium binding protein | EF hand family protein | NA | NA | 1 | No | No | No |
|  | TVAG_133030 | NADH dehydrogenase, Tvh-47 | Mitochondrial NADH dehydrogenase flavoprotein 1 | NA | NA | 3 | No | No | No |
|  | TVAG_212570 | Polymorphic outer membrane protein | Bifunctional inhibitor/lipid-transfer protein | NA | TM | 3 | Yes | Yes | No |
| **Other hydrolases** | TVAG_015160 | AP endonuclease | DNase I-like | NA | NA | 2 | No | No | No |
|  | TVAG_388060 | Saplip1 | Saposin | SP | NA | 2 | No | No | No |
|  | TVAG_164890 | Acetyl-CoA hydrolase | Acetyl-CoA hydrolase Ach1 | NA | NA | 2 | No | No | No |
| **Proteases** | TVAG_367130 | Clan MA, family M8, leishmanolysin-like metallopeptidase, GP63 | Peptidase M8, leishmanolysin | SP | TM | 2 | Yes | Yes | No |
|  | TVAG_034140 | Clan CA, family C1, papain-like cysteine peptidase | Peptidase C1A | NA | NA | 3 | Yes | No | Yes |
|  | TVAG_336300 | Clan AA, family A1, cathepsin D-like aspartic peptidase | Acid Protease | NA | NA | 3 | No | No | Yes |

NA = not available; SP = signal peptide; TBSR = *Trichomonas* beta-sandwich repeat-protein;
TM = transmembrane domain.

Phagocytosis is a typical mechanism by which TV acquires nutrients. Upon contact with various types of cells, including bacteria, leukocytes, *Saccharomyces cerevisiae*, and epithelial cells, TV cells form actin-driven pseudopodial extensions to surround the target cell, which are internalized and digested in lysosomes (6, 8, 9, 60). However, in our experiments, we observed that TV phagocytosed LJ without pseudopodia formation via a narrow opening in the plasma membrane, through which the LJ cell was suctioned into the cell. Only five minutes were sufficient to observe LJ inside TV by this process. Pseudopodia-independent phagocytosis by TV has been previously observed with the vaginal isolate of Döderlein’s bacteria and yeast (6, 9). This phenomenon is reminiscent of the sinking phagocytosis observed in macrophages and neutrophils (30–32). The molecular mechanism of sinking is not well understood, but it is different from classical phagocytosis and dependent on the stimulation of different receptor-opsonin combinations, with particular involvement of complement receptors in leukocytes (32). It is currently unclear whether pseudopodia-independent phagocytosis observed in TV and sinking relies on homologous mechanisms. Notably, we observed that TV exhibits significantly reduced phagocytic activity toward heat-inactivated LJ compared to viable bacteria. LJ is a non-motile bacterium that actively secretes various components, including surface-associated proteins with biosurfactant activity (61) and extracellular vesicles that influence cytoadherence (62). Identifying the native LJ-derived factors that mediate TV–LJ interactions will be essential for elucidating the molecular basis of pseudopodia-independent phagocytosis.

## Conclusions

A low abundance or complete absence of protective lactobacilli during acute trichomoniasis is a well-documented phenomenon (12, 13, 15, 63). However, the critical question remains whether alterations in the lactobacilli community occur prior to TV infection or if TV actively contributes to the depletion of these protective bacteria. Our results demonstrate that TV efficiently phagocytoses one of the dominant *Lactobacillus* species, LJ, through a rapid, pseudopodia-independent mechanism. Moreover, interaction with LJ stimulates the secretion of TvIS proteins, notably the TvGH25 lysozyme, linking secretory response to targeted antimicrobial activity against this *Lactobacillus* species. These findings contribute to a growing body of evidence suggesting that TV may actively participate in the displacement of healthy vaginal lactobacilli. Importantly, this insight identifies novel molecular targets for therapeutic strategies aimed at preserving the vaginal microbiome and enhancing microbial resilience during trichomoniasis.

## Materials and Methods

### Cell cultivation

The *T. vaginalis* strain TV17-48 (64) was cultured as described (65). For transfectant selection, the culture medium was supplemented with 200 µg/ml geneticin G418 or 40 μg/ml puromycin (10). *L. jensenii* CECT 4306 (66) was cultured as described (67). Agar in a final concentration of 0.9% (m/v) was used for cultivation on plates. *Candida albicans* SC5314 was provided by Robert Hirt (Newcastle University, United Kingdom) and cultured as described (68).

### Gene cloning and *T. vaginalis* transfection

The genes encoding TvGH25 (TVAG_088640) and Hybrid Cluster Protein 1 (HCP1, TVAG_206500) were cloned into the vectors pTagVag-V5-Pur (10) or pTagVag-HA-Neo (69), respectively, fusing the proteins with a V5 tag or haemagglutinin (HA) tag at their C-terminus. Both genes were overexpressed using the strong promoter of α-subunit succinyl-CoA synthetase (SCS)(42) and transfected as described (70). Details are given in *SI Appendix, Materials and Methods*, and Table S2.

### *Trichomonas*-*Lactobacillus* co-incubations

For all co-incubations, overnight cultures of TV were harvested by centrifugation (10 min at 1,000 x g) and washed in isotonic Doran’s medium (71) supplemented with 15 mM maltose. LJ was grown to an OD600 of 0.4 (logarithmic phase), harvested by centrifugation (5 min at 5,000 x g), and washed as TV. Then, both TV and LJ were resuspended and combined in either TYM or Doran’s medium to a final concentration of 10^6^/ml (TV) and 10^7^/ml (LJ), respectively, resulting in a TV-to-LJ ratio of 1:10. Subsequent incubation was performed at 37 °C for time intervals detailed in the respective experimental descriptions.

### Experimental design and statistical rationale

Each experiment was performed in three independent biological replicates. Unless specified otherwise, one-way ANOVA was applied on the composite data for statistical analyses, and a t-test to compare the secretion scores.

### ImageStream flow cytometric analysis of *T. vaginalis* phagocytic activity

Lactobacilli were stained with BacLight™ Red (10 nM final concentration; Thermo Fisher Scientific) at 37°C for 30 min and then washed with Doran’s medium. TV was harvested and washed, mixed with stained lactobacilli, and co-incubated in Doran’s medium for 5, 15, and 30 min at 37 °C. Co-incubation at 15°C served as a control. Then, the samples were analyzed with the Amnis® ImageStream®X Mk II Flow Cytometer (Cytek, USA) until a minimum of 1.2 x 10^3^ TV cells per sample were detected. Images were processed using the INSPIRE^TM^ software version 201.1.0.765 (Cytek, USA), followed by the analysis with Python software version 3.10.17 with the libraries CellPose v3.1.1.1, Scikit-image v0.25.0, SciPy v1.15.3, and Scikit-learn v1.6.1.

### *L. jensenii* growth following co-incubation

TV and LJ were harvested, washed, and co-incubated in TYM as above (TV-to-LJ ratio of 1:10) for 10 to 300 min. Aliquots were taken after each time point, ten-fold serial dilutions were prepared for final dilution factors ranging from 1:10^2^ to 1:10^4^, plated on agar plates, and incubated overnight at 37 °C. As a control, LJ was incubated without TV. Images of the plates were taken using the Amersham Imager 600 (GE Healthcare Life Sciences) and edited in ImageJ/Fiji version 1.54p (72) to increase brightness and contrast of the colony-forming units (CFUs). Plates with optimal dilution that resulted in clearly separated CFUs were selected for counting, and CFU/ml was calculated by multiplying the colony count by the reciprocal of the dilution factor and dividing by the plated volume. CFUs were counted using OpenCFU version 3.9 (73) with threshold set to ‘inverted’ and both threshold and minimum radius set to the value of 1.

### Fluorescence microscopy of phagocytosis assays

To confirm phagocytosis of LJ by TV via microscopy, TV cells overexpressing HA-tagged Rab7 were used for the co-incubation to visualize the endolysosomal system (29). After co-incubation for 30 min, the cells were processed as described (74) using mouse monoclonal α-HA antibody (Exbio) and donkey α-mouse antibody conjugated to Alexa Fluor 488 (Thermo Fisher). Samples were mounted using DAPI-containing mounting medium (Vectashield) to visualize LJ. Details are given in *SI Appendix, Materials and Methods*.

### Transmission electron microscopy (TEM)

TV and LJ were co-incubated in TYM for 5 min and 15 min as above, fixed in 2.5% glutaraldehyde, and processed as described (75). Details are given in *SI Appendix, Materials and Methods*.

### Scanning electron microscopy (SEM)

To analyze surface structures involved in phagocytosis, TV and LJ cells were co-incubated in TYM for 5 min and 15 min as above. As a control, TV was co-incubated with *C. albicans* for 15, 60, and 120 min as described (6). Then, the cells were prefixed with 1% formaldehyde, washed with PBS, and placed on a cover glass. Next, they were fixed with 2.5% glutaraldehyde in 0.1 M cacodylate buffer (pH 7.2). The samples were washed with 0.1 M cacodylate buffer (pH 7.2), post-fixed with 2% OsO_4_ in cacodylate buffer (pH 7.2), washed again with distilled water, dehydrated using an ascending series of ethanol and acetone, and dried with CO_2_ in a critical point dryer. Finally, the samples were coated with a thin gold/platinum layer using an ion sputter coater. The finalized samples were observed with the FEI Helios NanoLab 660 G3 UC microscope (Thermo Fisher Scientific).

### Incubation of *T. vaginalis* with inactive and viable *L. jensenii*

To prepare heat-inactivated LJ (HiLJ), LJ cells were harvested as above, resuspended in 100 µl Doran’s medium, and inactivated at 95°C for 30 min. To confirm the absence of viable LJ cells, an aliquot was plated on an agar plate and incubated overnight at 37 °C. Next, viable LJ (vLJ) were harvested, and both HiLJ and vLJ were stained with BacLight™ Red and washed as above. TV was co-incubated for 30 min at 37°C with either vLJ or HiLJ in a ratio of 1:10 (TV to lactobacilli). The samples were analyzed with the Amnis® ImageStream®X Mk II Flow Cytometer (Cytek, USA).

### Secretome analysis by mass spectrometry (MS)

TV and LJ were harvested and washed twice as before and then mixed in Doran’s to a final concentration of 10^6^ cells/ml (TV) and 10^7^ cells/ml (LJ), respectively, and 15 ml of the suspensions were incubated anaerobically in culture flasks (25 cm², plug cap, SPL Life Sciences) for 10, 30, and 60 min at 37 °C. Control cells were incubated for 60 min on ice with or without LJ. After incubation, the supernatants were collected and remaining cells were removed by centrifugation at 1,000 × g for 5 min at 4 °C. Next, the supernatants were centrifuged at 10,000 × g for 10 min to remove cell debris, filtered through a 0.22 μm filter (Roth), and centrifuged at 100,000 × g for 75 min to remove microvesicles (2). The proteins in the final supernatants were precipitated with trichloroacetic acid as described (76) and stored at −80 °C. Label-free quantitative (LFQ) mass spectrometry (MS) analysis was performed as described previously (10, 77). Each biological sample was analyzed by MS in three technical replicates. To ensure TV cell integrity during the incubations, at each time point, we determined the extracellular activity of the cytosolic enzyme NADH oxidase in the cell suspensions (78). The maximum acceptable extracellular enzyme activity was set to 5% of the total activity.

### MS data acquisition

Tryptic peptides were injected on a nano reverse-phase liquid chromatograph (UltiMate 3000 RSLC, Thermo Fisher Scientific) coupled with MS (nanoLC-MS) using an Orbitrap Fusion Tribrid mass spectrometer (Thermo Fisher Scientific) as described previously (10). Details are given in *SI Appendix, Materials and Methods*.

### Analysis of MS data

Raw data were processed, and the secreted proteins initially filtered as described (10, 79, 80, and *SI Appendix, Materials and Methods*). The resulting lists of proteins from the axenic culture and co-culture were combined and compared. The z-score normalized LFQs from the time points 30 and 60 minutes were clustered and visualized using the R package pheatmap (81). The distances between the proteins were calculated using Pearson correlation, and Ward’s method was applied to cluster the proteins according to distances. The differences between the LFQs of the axenic culture and co-culture from the same time points were calculated, and the significance of the differences in each group was determined by the Kruskal-Wallis test. The significance of the secretion scores between the axenic culture and co-culture in each group was determined by a classical t-test. Then, for each protein, the difference in secretion score between the axenic culture and the co-culture was calculated. The proteins of the induced secretome were filtered based on the following criteria: (i) the protein’s secretion score difference was ≥ 0.5, (ii) the LFQ difference of the protein’s group increased significantly over time, and (iii) the secretion score of the protein’s group was significantly higher in the co-culture than in the axenic control. Based on these criteria, a heatmap of the interactome was constructed.

### Bioinformatics

For each protein of the induced secretome, signal peptides were predicted using SignalP-5.0 and transmembrane domains using DeepTMHMM version 1.0.42. Conserved domains were predicted using the database InterPro version 104.0 (47,677 entries). Identified proteins were annotated and categorized based on TrichDB annotations.

### RNA isolation and Real-time quantitative PCR

Total RNA was isolated according to the manufacturer’s protocol using the High Pure RNA Isolation Kit (Roche). Real-time quantitative PCR (RT-qPCR) was performed according to the manufacturer’s protocol in the Rotor-Gene 3000 RT-qPCR cycler (Qiagen) and evaluated using standard methods described in *SI Appendix, Materials and Methods*.

### Preparation of recombinant TvGH25 and mutTvGH25

TvGH25 was cloned into the vector pET-42b(+), fusing the protein with a C-terminal polyhistidine (His) tag. A mutated version of the enzyme, mutTvGH25, with two point mutations in the active site (Asp100Ala, Glu102Gln), was prepared by fusing two overlapping PCR-amplified fragments coding the mutated amino acids. MutTvGH25 was cloned into the same vector as TvGH25. His-tagged TvGH25 and mutTvGH25 were expressed in and purified from *E. coli* BL-21 (DE3) as described (82). Details are given in *SI Appendix, Materials and Methods*, and Table S2.

### Muramidase activity of TvGH25

PG from *E. coli* BW25113Δ6LDT and *L. gasseri* ATCC 9857 were prepared as described (40, 84). The substrates were incubated with TvGH25 or mutTvGH25 (5 µM) in 20 mM sodium phosphate pH 4.8 for 18 h at 37°C using a thermal shaker set at 900 rpm. Control PG samples were incubated with 4 µM (PG from *E. coli*) and 2 µM (PG from *L. gasseri*) of the muramidase cellosyl (Hoechst, Frankfurt) or without enzyme. The reactions were stopped by heating the samples for 10 min in a dry heat block set at 100°C. Samples were centrifuged at 17,000 × g for 15 min, and the supernatant was retrieved. Samples were reduced by adding one sample volume of 0.7 M sodium borate (pH 9.0) with 10 mg/ml sodium borohydride and incubating for 30 min at room temperature. Samples were acidified to pH 3.5-4.5 with 20% phosphoric acid and separated by high-performance liquid chromatography (HPLC) using published methods for *E. coli* (83) and *L. gasseri* (84) samples.

### TvGH25 localization by expansion fluorescence microscopy

The V5-tagged TvGH25 transfectants were prefixed with 1% formaldehyde and processed as described (85). The cells were stained using mouse monoclonal anti-V5 antibody (ab206558, ABCAM) and donkey anti-mouse antibody conjugated to Alexa Fluor 488 (Thermo Fisher Scientific). The fluorescent probe ATTO 550 NHS ester (Thermo Fisher Scientific) was used to label protein-rich cell structures. Finalized samples were observed with a Nikon CSU-W1 spinning disc confocal microscope (Nikon Healthcare). Images were deconvolved using Huygens Professional version 19.10 (Scientific Volume Imaging) and further processed using ImageJ/Fiji software (72) and the Imaris 9.7.2 Package for Cell Biologists (Bitplane AG).

## Acknowledgements

The authors thank Luis Bermudez-Humaran (Institut national de la recherche agronomique, France) and Robert Hirt (Newcastle University, United Kingdom) for providing *L. jensenii* and *C. albicans* isolates, respectively. The authors acknowledge Matyáš Šíma from the Flow Cytometry unit at IMG and Zuzana Čočková from the Imaging Methods Core Facility at BIOCEV for their technical support in flow cytometric data acquisition and image cytometry analysis, respectively. The authors acknowledge the OMICS Proteomics at BIOCEV.

## References

1. World Health Organization, Report on global sexually transmitted infection surveillance 2018 (World Health Organization, 2018).

2. O. Twu, et al., *Trichomonas vaginalis* homolog of macrophage migration inhibitory factor induces prostate cell growth, invasiveness, and inflammatory responses. Proc. Natl. Acad. Sci. U. S. A. 111, 8179–8184 (2014).

3. P. Kissinger, *Trichomonas vaginalis*: A review of epidemiologic, clinical and treatment issues. BMC Infect. Dis. 15 (2015).

4. D. Petrin, K. Delgaty, R. Bhatt, G. Garber, Clinical and microbiological aspects of *Trichomonas vaginalis*. Clin. Microbiol. Rev. 11, 300–317 (1998).

5. P. Kissinger, A. Adamski, Trichomoniasis and HIV interactions: A review. Sex. Transm. Infect. 89, 426–433 (2013).

6. A. Pereira-Neves, M. Benchimol, Phagocytosis by *Trichomonas vaginalis*: new insights. Biol. Cell 99, 87–101 (2007).

7. M. Benchimol, I. De Andrade Rosa, R. Da Silva Fontes, Â. J. Burla Dias, *Trichomonas* adhere and phagocytose sperm cells: Adhesion seems to be a prominent stage during interaction. Parasitol. Res. 102, 597–604 (2008).

8. V. Midlej, M. Benchimol, *Trichomonas vaginalis* kills and eats -Evidence for phagocytic activity as a cytopathic effect. Parasitology 137, 65–76 (2010).

9. J. G. Rendón-Maldonado, M. Espinosa-Cantellano, A. González-Robles, A. Martínez-Palomo, *Trichomonas vaginalis*: In vitro phagocytosis of lactobacilli, vaginal epithelial cells, leukocytes, and erythrocytes. Exp. Parasitol. 89, 241–250 (1998).

10. J. Štáfková, et al., Dynamic secretome of *Trichomonas vaginalis*: Case study of β-amylases. Mol. Cell. Proteomics 17, 304–320 (2018).

11. J. Pinheiro, et al., The Protozoan *Trichomonas vaginalis* Targets Bacteria with Laterally Acquired NlpC/P60 Peptidoglycan Hydrolases. mBio 9 (2018).

12. R. M. Brotman, et al., Association between *Trichomonas vaginalis* and vaginal bacterial community composition among reproductive-age women. Sex. Transm. Dis. 39, 807 (2012).

13. S. F. Chiu, et al., Vaginal Microbiota of the Sexually Transmitted Infections Caused by *Chlamydia trachomatis* and *Trichomonas vaginalis* in Women with Vaginitis in Taiwan. Microorganisms 9 (2021).

14. J. Ravel, et al., Vaginal microbiome of reproductive-age women. Proc. Natl. Acad. Sci. U. S. A. 108, 4680–4687 (2011).

15. N. Kalia, J. Singh, M. Kaur, Microbiota in vaginal health and pathogenesis of recurrent vulvovaginal infections: a critical review. Ann. Clin. Microbiol. Antimicrob. 19, 5 (2020).

16. J. Leizer, et al., 447: Large differences in the composition of vaginal secretions in pregnant women in the presence of *Lactobacillus crispatus* and *L. iners*. Am. J. Obstet. Gynecol. 216, S264 (2017).

17. N. Phukan, T. Parsamand, A. Brooks, T. Nguyen, A. Simoes-Barbosa, The adherence of *Trichomonas vaginalis* to host ectocervical cells is influenced by lactobacilli. Sex. Transm. Infect. 89, 455–459 (2013).

18. N. Phukan, A. E. S. Brooks, A. Simoes-Barbosa, A Cell Surface Aggregation-Promoting Factor from *Lactobacillus gasseri* Contributes to Inhibition of *Trichomonas vaginalis* Adhesion to Human Vaginal Ectocervical Cells. Infect. Immun. 86, e00907–17 (2018).

19. B. Pradines, S. Domenichini, V. Lievin-Le Moal, Adherent Bacteria and Parasiticidal Secretion Products of Human Cervicovaginal Microbiota-Associated *Lactobacillus gasseri* Confer Non-Identical Cell Protection against *Trichomonas vaginalis*-Induced Cell Detachment. Pharmaceuticals 15, 1350 (2022).

20. W. J. Y. Chee, S. Y. Chew, L. T. L. Than, Vaginal microbiota and the potential of *Lactobacillus* derivatives in maintaining vaginal health. Microb. Cell Factories 19, 203 (2020).

21. A. S. Hinderfeld, N. Phukan, A.-K. Bär, A. M. Roberton, A. Simoes-Barbosa, Cooperative Interactions between *Trichomonas vaginalis* and Associated Bacteria Enhance Paracellular Permeability of the Cervicovaginal Epithelium by Dysregulating Tight Junctions. Infect. Immun. 87, e00141–19 (2019).

22. U. S. Fatić, L. Lekić, G. Malesevic, V. S. Stanković, N. P. Markota, Effectiveness of *Lactobacillus jensenii* in treating vaginal and urogenital infections: Age-related differences in response and immune function. MEDIS – Int. J. Med. Sci. Res. 4, 27–31 (2025).

23. V. G. Nair, et al., Human vaginal Lactobacillus Jensenii -derived (-)-Terpinen-4-ol restores antibiotic sensitivity by inhibiting efflux pumps in drug resistant E. coli and K. pneumoniae. Sci. Rep. 15, 31823 (2025).

24. Q. Zhang, et al., Targeting vaginal dysbiosis: prospects for the application of live biotherapeutics products. Front. Microbiol. 17, 1749581.

25. L. Qin, et al., Population-level analyses identify host and environmental variables influencing the vaginal microbiome. Signal Transduct. Target. Ther. 10, 64 (2025).

26. N. Zheng, R. Guo, J. Wang, W. Zhou, Z. Ling, Contribution of *Lactobacillus iners* to Vaginal Health and Diseases: A Systematic Review. Front. Cell. Infect. Microbiol. 11, 792787 (2021).

27. M. Vaneechoutte, *Lactobacillus iners*, the unusual suspect. Res. Microbiol. 168, 826–836 (2017).

28. M. I. Petrova, E. Lievens, S. Malik, N. Imholz, S. Lebeer, *Lactobacillus* species as biomarkers and agents that can promote various aspects of vaginal health. Front. Physiol. 6, 81 (2015).

29. N. Zimmann, et al., Proteomic Analysis of *Trichomonas vaginalis* Phagolysosome, Lysosomal Targeting, and Unconventional Secretion of Cysteine Peptidases. Mol. Cell. Proteomics 21, 100174 (2022).

30. M. B. Hallett, Phagocytosis of optically-trapped particles: delivery of the pure phagocytic signal. Cell Res. 16, 852–854 (2006).

31. E. Gagnon, et al., Endoplasmic reticulum-mediated phagocytosis is a mechanism of entry into macrophages. Cell 110, 119–131 (2002).

32. S. Walbaum, et al., Complement receptor 3 mediates both sinking phagocytosis and phagocytic cup formation via distinct mechanisms. J. Biol. Chem. 296, 100256 (2021).

33. J. M. Halbleib, W. J. Nelson, Cadherins in development: cell adhesion, sorting, and tissue morphogenesis. Genes Dev. 20, 3199–3214 (2006).

34. O. Klezovitch, V. Vasioukhin, Cadherin signaling: keeping cells in touch. [Preprint] (2015). Available at: https://f1000research.com/articles/4-550 [Accessed 27 June 2025].

35. T. Pachano, et al., Epigenetics regulates transcription and pathogenesis in the parasite *Trichomonas vaginalis*. Cell. Microbiol. 19 (2017).

36. R. Herbst, F. Marciano-Cabral, M. Leippe, Antimicrobial and pore-forming peptides of free-living and potentially highly pathogenic *Naegleria fowleri* are released from the same precursor molecule. J. Biol. Chem. 279, 25955–25958 (2004).

37. M. Leippe, J. Andrä, R. Nickel, E. Tannich, H. J. Müller-Eberhard, Amoebapores, a family of membranolytic peptides from cytoplasmic granules of *Entamoeba histolytica*: isolation, primary structure, and pore formation in bacterial cytoplasmic membranes. Mol. Microbiol. 14, 895– 904 (1994).

38. L. A. Mpeyako, et al., Comparative genomics between *Trichomonas tenax* and *Trichomonas vaginalis*: CAZymes and candidate virulence factors. Front. Microbiol. 15, 1437572 (2024).

39. J. A. Metcalf, L. J. Funkhouser-Jones, K. Brileya, A.-L. Reysenbach, S. R. Bordenstein, Antibacterial gene transfer across the tree of life. eLife 3, e04266 (2014).

40. E. Kuru, et al., Fluorescent D-amino-acids reveal bi-cellular cell wall modifications important for *Bdellovibrio bacteriovorus* predation. Nat. Microbiol. 2, 1648–1657 (2017).

41. A. Rau, T. Hogg, R. Marquardt, R. Hilgenfeld, A new lysozyme fold. Crystal structure of the muramidase from *Streptomyces coelicolor* at 1.65 A resolution. J. Biol. Chem. 276, 31994–31999 (2001).

42. P. Rada, et al., N-Terminal Presequence-Independent Import of Phosphofructokinase into Hydrogenosomes of *Trichomonas vaginalis*. Eukaryot. Cell 14, 1264–1275 (2015).

43. S. A. Ragland, A. K. Criss, From bacterial killing to immune modulation: Recent insights into the functions of lysozyme. PLOS Pathog. 13, e1006512 (2017).

44. W. Vollmer, B. Joris, P. Charlier, S. Foster, Bacterial peptidoglycan (murein) hydrolases. FEMS Microbiol. Rev. 32, 259–286 (2008).

45. M. Benchimol, C. Batista, W. De Souza, Fibronectin-and laminin-mediated endocytic activity in the parasitic protozoa *Trichomonas vaginalis* and *Tritrichomonas foetus*. J. Submicrosc. Cytol. Pathol. 22, 39–45 (1990).

46. A. Lanfredi-Rangel, M. Attias, T. M. de Carvalho, W. M. Kattenbach, W. De Souza, The peripheral vesicles of trophozoites of the primitive protozoan *Giardia lamblia* may correspond to early and late endosomes and to lysosomes. J. Struct. Biol. 123, 225–235 (1998).

47. M. Benchimol, W. de Souza, Endocytosis in anaerobic parasitic protists. Mem. Inst. Oswaldo Cruz 119, e240058.

48. R. Santos, et al., Combined nanometric and phylogenetic analysis of unique endocytic compartments in *Giardia lamblia* sheds light on the evolution of endocytosis in Metamonada. BMC Biol. 20, 206 (2022).

49. M. J. Barnett, et al., NlpC/P60 peptidoglycan hydrolases of *Trichomonas vaginalis* have complementary activities that empower the protozoan to control host-protective lactobacilli. PLOS Pathog. 19, e1011563 (2023).

50. C. Juliano, P. Cappuccinelli, A. Mattana, In vitro phagocytic interaction between *Trichomonas vaginalis* isolates and bacteria. Eur. J. Clin. Microbiol. Infect. Dis. 10, 497–502 (1991).

51. G. Lustig, C. M. Ryan, W. E. Secor, P. J. Johnson, *Trichomonas vaginalis* contact-dependent cytolysis of epithelial cells. Infect. Immun. 81, 1411–1419 (2013).

52. R. Arroyo, et al., *Trichomonas vaginalis* Cysteine Proteinases: Iron Response in Gene Expression and Proteolytic Activity. BioMed Res. Int. 2015 (2015).

53. L. I. Quintas-Granados, et al., TvMP50 is an immunogenic metalloproteinase during male trichomoniasis. Mol. Cell. Proteomics MCP 12, 1953–1964 (2013).

54. S. Nakjang, D. A. Ndeh, A. Wipat, D. N. Bolam, R. P. Hirt, A Novel Extracellular Metallopeptidase Domain Shared by Animal Host-Associated Mutualistic and Pathogenic Microbes. PLoS ONE 7, e30287 (2012).

55. M. Leippe, R. Herbst, Ancient weapons for attack and defense: the pore-forming polypeptides of pathogenic enteric and free-living amoeboid protozoa. J. Eukaryot. Microbiol. 51, 516–521 (2004).

56. N. Diaz, et al., Production and Functional Characterization of a Recombinant Predicted Pore-Forming Protein (TVSAPLIP12) of *Trichomonas vaginalis* in *Nicotiana benthamiana* Plants. Front. Cell. Infect. Microbiol. 10, 581066 (2020).

57. N. De Miguel, et al., Proteome analysis of the surface of *Trichomonas vaginalis* reveals novel proteins and strain-dependent differential expression. Mol. Cell. Proteomics 9, 1554–1566 (2010).

58. Y.-P. Chen, A. M. Riestra, A. K. Rai, P. J. Johnson, A Novel Cadherin-like Protein Mediates Adherence to and Killing of Host Cells by the Parasite *Trichomonas vaginalis*. mBio 10, e00720–19 (2019).

59. A. M. Riestra, et al., A *Trichomonas vaginalis* Rhomboid Protease and Its Substrate Modulate Parasite Attachment and Cytolysis of Host Cells. PLoS Pathog. 11 (2015).

60. P. Francioli, H. Shio, R. B. Roberts, M. Müller, Phagocytosis and killing of *Neisseria gonorrhoeae* by *Trichomonas vaginalis*. J. Infect. Dis. 147, 87–94 (1983).

61. R. R. Spurbeck, C. G. Arvidson, Lactobacillus jensenii Surface-Associated Proteins Inhibit Neisseria gonorrhoeae Adherence to Epithelial Cells. Infect. Immun. 78, 3103–3111 (2010).

62. A. Artuyants, J. Hong, P. Dauros-Singorenko, A. Phillips, A. Simoes-Barbosa, *Lactobacillus gasseri* and *Gardnerella vaginalis* produce extracellular vesicles that contribute to the function of the vaginal microbiome and modulate host-*Trichomonas vaginalis* interactions. Mol. Microbiol. 122, 357–371 (2024).

63. V. Margarita, P. L. Fiori, P. Rappelli, Impact of Symbiosis Between *Trichomonas vaginalis* and *Mycoplasma hominis* on Vaginal Dysbiosis: A Mini Review. Front. Cell. Infect. Microbiol. 10, 179 (2020).

64. J. Kulda, M. Vojtechovska, J. Tachezy, P. Demes, E. Kunzová, Metronidazole resistance of *Trichomonas vaginalis* as a cause of treatment failure in trichomoniasis. A case report. Br. J. Vener. Dis. 58, 394–399 (1982).

65. L. S. Diamond, The establishment of various trichomonads of animals and man in axenic cultures. J. Parasitol. 43, 488–490 (1957).

66. R. Martín, et al., Effect of iron on the probiotic properties of the vaginal isolate *Lactobacillus jensenii* CECT 4306. Microbiol. Read. 708–18 (2015).

67. J. de Man, M. Rogosa, M. E. Sharpe, A Medium for the Cultivation of Lactobacilli. J. Appl. Bacteriol. 23, 130–135 (1960).

68. YPD media. Cold Spring Harb. Protoc. 2010, pdb.rec12315 (2010).

69. I. Hrdý, et al., *Trichomonas* hydrogenosomes contain the NADH dehydrogenase module of mitochondrial complex I. Nature 432, 618–622 (2004).

70. B. D. Janssen, et al., CRISPR/Cas9-mediated gene modification and gene knock out in the human-infective parasite *Trichomonas vaginalis*. Sci. Rep. 8 (2018).

71. D. J. Doran, Studies on Trichomonads. III. Inhibitors, Acid Production, and Substrate Utilization by 4 Strains of *Tritrichomonas foetus*. J. Protozool. 6, 177–182 (1959).

72. J. Schindelin, et al., Fiji: An open-source platform for biological-image analysis. Nat. Methods 9, 676–682 (2012).

73. Q. Geissmann, OpenCFU, a New Free and Open-Source Software to Count Cell Colonies and Other Circular Objects. PLoS ONE 8 (2013).

74. D. J. Woessner, S. C. Dawson, The *Giardia* Median Body Protein Is a Ventral Disc Protein That Is Critical for Maintaining a Domed Disc Conformation during Attachment. Eukaryot. Cell 11, 292 (2012).

75. N. C. Beltrán, et al., Iron-Induced Changes in the Proteome of *Trichomonas vaginalis* Hydrogenosomes. PLoS ONE 8, 65148 (2013).

76. V. Méchin, C. Damerval, M. Zivy, Total protein extraction with TCA-acetone. Methods Mol. Biol. Clifton NJ 355, 1–8 (2007).

77. T. Masuda, M. Tomita, Y. Ishihama, Phase transfer surfactant-aided trypsin digestion for membrane proteome analysis. J. Proteome Res. 7, 731–740 (2008).

78. D. Linstead, S. Bradley, The purification and properties of two soluble reduced nicotinamide: Acceptor oxidoreductases from *Trichomonas vaginalis*. Mol. Biochem. Parasitol. 27, 125–133 (1988).

79. J. Cox, et al., Accurate proteome-wide label-free quantification by delayed normalization and maximal peptide ratio extraction, termed MaxLFQ. Mol. Cell. Proteomics 13, 2513–2526 (2014).

80. J. Alvarez-Jarreta, et al., VEuPathDB: the eukaryotic pathogen, vector and host bioinformatics resource center in 2023. Nucleic Acids Res. 52, D808–D816 (2024).

81. R. Kolde, pheatmap: Pretty Heatmaps. (2018).

82. E. Nývltová, T. Smutná, J. Tachezy, I. Hrdý, OsmC and incomplete glycine decarboxylase complex mediate reductive detoxification of peroxides in hydrogenosomes of *Trichomonas vaginalis*. Mol. Biochem. Parasitol. 206, 29–38 (2016).

83. B. Glauner, Separation and quantification of muropeptides with high-performance liquid chromatography. Anal. Biochem. 172, 451–464 (1988).

84. N. K. Bui, et al., Isolation and analysis of cell wall components from Streptococcus pneumoniae. Anal. Biochem. 421, 657–666 (2012).

85. P. Gorilak, et al., Expansion microscopy facilitates quantitative super-resolution studies of cytoskeletal structures in kinetoplastid parasites. Open Biol. 11, 210131 (2021).

