## Supporting Information for "*Trichomonas vaginalis* targets *Lactobacillus jensenii* via pseudopodia-independent phagocytosis and secreted lysozyme TvGH25" (from another version of this preprint: version 2, 2026-07-31)

**This PDF file includes:**

Materials and Methods
Figures S1 to S5
Tables S1 to S2
Legends for Datasets S1 to S2
SI References

**Other supporting materials for this manuscript include the following:**

Datasets S1 to S2

### Materials and Methods

Gene cloning and *T. vaginalis* transfection

The plasmids were transfected into trichomonads by nucleofection using the Human T Cell Nucleofector Kit (Lonza). Briefly, 6 x $10^6$ cells in the logarithmic growth phase were harvested by centrifugation (10 min at 1,000 x g) and gently resuspended in 82 µl Nucleofector Solution and 18 µl Supplement 1. Then, 10 µg of plasmid was added, and the mixture was incubated at room temperature for 10 min. The manufacturer's program U.033 was used for transfection (1).

ImageStream flow cytometric analysis of *T. vaginalis* phagocytic activity

In INSPIRE, appropriate gating was applied to obtain *T. vaginalis* populations using image contrast and area. Further gating was done using the area and intensity of the BacLight Red signal. Exported images consisted of three channels (brightfield, BacLight Red, side scatter) and were further processed in the Python software version 3.10.17 using the libraries CellPose v3.1.1.1, Scikit-image v0.25.0, SciPy v1.15.3, and Scikit-learn v1.6.1. Dead TV cells were excluded based on the side scatter, and doublets were excluded by masks. Automated segmentation was set up to analyze only single-TV cell events. Classification accuracy reached 98.08% on a training dataset comprising 1,557 images. For final image analysis, the IDEAS® software version 6.4 was used.

Fluorescence microscopy of phagocytosis assays

Slides were observed with a Leica TCS SP8 confocal laser scanning microscope (Leica Microsystems). Images were processed using Huygens Professional version 19.10 (Scientific Volume Imaging) and further processed using ImageJ/Fiji version 1.54p (2) and the Imaris 9.7.2 Package for Cell Biologists (Bitplane AG).

Transmission electron microscopy

After co-incubation, the cells were centrifuged at 3 000 × g for 10 min and fixed overnight at 4°C in 2.5% glutaraldehyde and 5 mM $CaCl_2$ in 0.1 M cacodylate buffer (pH 7.2). Subsequently, the cells were postfixed for 15 min at 4°C in 0.1 M cacodylate buffer containing 1.6% ferricyanide, 10 mM $CaCl_2$, and 2% $OsO_4$. Fixed samples were dehydrated using an ascending series of ethanol and acetone and embedded in the EMBed 812 epoxy resin (Electron Microscopy Sciences, Hatfield, PA, USA). Ultrathin sections were stained with uranyl acetate and observed using a JEOL JEM-1011 microscope.

MS data acquisition

Peptides were loaded onto an Acclaim PepMap300 trap column (300 µm × 5 mm) packed with C18 (5 µm, 300 Å) in loading buffer (0.1% trifluoroacetic acid in 2% acetonitrile) for 4 min at a flow rate of 15 µl/min and then separated on an EASY-Spray column (75 µm × 50 cm) packed with C18 (2 µm, 100 Å, Thermo Fisher Scientific) at a flow rate of 300 nl/min. Mobile phase A (0.1% formic acid in water) and mobile phase B (0.1% formic acid in acetonitrile) were used to establish a 60-min gradient from 4% to 35% B. Eluted peptides were ionized by electrospray. The full MS spectrum (350–1400 m/z range) was acquired at a resolution of 120,000 at m/z 200 and a maximum ion accumulation time of 100 ms. Dynamic exclusion was set to 60 s. Higher-energy collisional dissociation (HCD) MS/MS spectra were acquired in iontrap in rapid mode, and the normalized collision energy was set to 30% with a maximum ion accumulation time of 35 ms. An isolation width of 1.6 m/z units was used for MS 2.

Analysis of MS data

Raw data were processed using MaxQuant version 2.3.0.0 (Max-Planck-Institute of Biochemistry) (3). Searches were performed using the *T. vaginalis* database 'tvag_eupath_and_uniprot_20191115' from EUpathDB (4) (97,729 entries), and *L. jensenii* database from UniProt (4,897 entries). Trypsin cleaving specificity was used to generate the peptides, and two missed cleavages were allowed. The protein modifications were set as follows: cysteine (unimod nr: 39) as static, and methionine oxidation (unimod: 1384) and protein N-terminus acetylation (unimod: 1) as variable. The false discovery rates for peptides and proteins were set to 1%. Search mass tolerances were used in MaxQuant default settings for Orbitrap and Iontrap. The precursor ion mass tolerance in the initial search was 20 ppm, the tolerance in the main search was 4.5 ppm, and the fragment ion mass tolerance was 0.5 Da.

Secreted proteins were filtered using the following criteria: (i) the protein was identified by at least three peptides, (ii) the protein increased by at least 1 LFQ unit in comparison to the control trichomonads kept on ice, and (iii) the protein increased significantly over time. The significance of the increase was evaluated using the Friedman test, and the value of 0.1 was set as a cut-off. For each protein, the secretion score was calculated as previously described (5). The resulting lists of proteins from the axenic culture and co-culture were combined and compared. The z-score normalized LFQs from the time points 30 and 60 minutes were clustered and visualized using the R package pheatmap (6). The distances between the proteins were calculated using Pearson correlation, and Ward's method was applied to cluster the proteins according to distances. The differences between the LFQs of the axenic culture and co-culture from the same time points were calculated, and the significance of the differences in each group was determined by the Kruskal-Wallis test. The significance of the secretion scores between the axenic culture and co-culture in each group was determined by a classical t-test. Then, for each protein, the difference in secretion score between the axenic culture and the co-culture was calculated. The proteins of the induced secretome were filtered based on the following criteria: (i) the protein's secretion score difference was ≥ 0.5, (ii) the LFQ difference of the protein's group increased significantly over time, and (iii) the secretion score of the protein's group was significantly higher in the co-culture than in the axenic control. Based on these criteria, a heatmap of the interactome was constructed.

Preparation of recombinant TvGH25 and mutTvGH25 from *E. coli*

Bacterial cultures were grown in LB (Lennox) medium (Sigma-Aldrich) at 37 °C to $OD_{A600}$ = 0.8, then protein expression was induced by the addition of 0.5 mM isopropyl-β-d-1-thiogalactopyranoside (IPTG) for 4 hours. Samples were homogenized using a French Press High Pressure Cell (GlenMills, USA) at 23 kPSI, followed by centrifugation (120,000 × g for 35 min at 4 °C). The proteins were purified from the soluble fraction by affinity chromatography on a Ni-NTA agarose column under native conditions according to the manufacturer's protocol (Qiagen). The purified proteins were transferred to PBS (pH 7.2) and concentrated using an Amicon Ultra concentrator (Millipore; 10 kDa pore size).

RNA isolation and Real-time quantitative PCR

TV and LJ were co-incubated and harvested as above. Total RNA was isolated according to the manufacturer's protocol using the High Pure RNA Isolation Kit (Roche). Real-time quantitative PCR (RT-qPCR) was performed according to the manufacturer's protocol in the Rotor-Gene 3000 RT-qPCR cycler (Qiagen) using the Kapa SYBR FAST Universal One-step RT-qPCR kit (Sigma-Aldrich) with specific primers for TvGH25, TBSR1, and TBSR12 (Table S2). The expression of topoisomerase II (TVAG_038880) was used for the normalization of expression levels of the genes analyzed. The relative expression was determined using the $2^{-\Delta\Delta Ct}$ method (7).

### Supplementary Figures

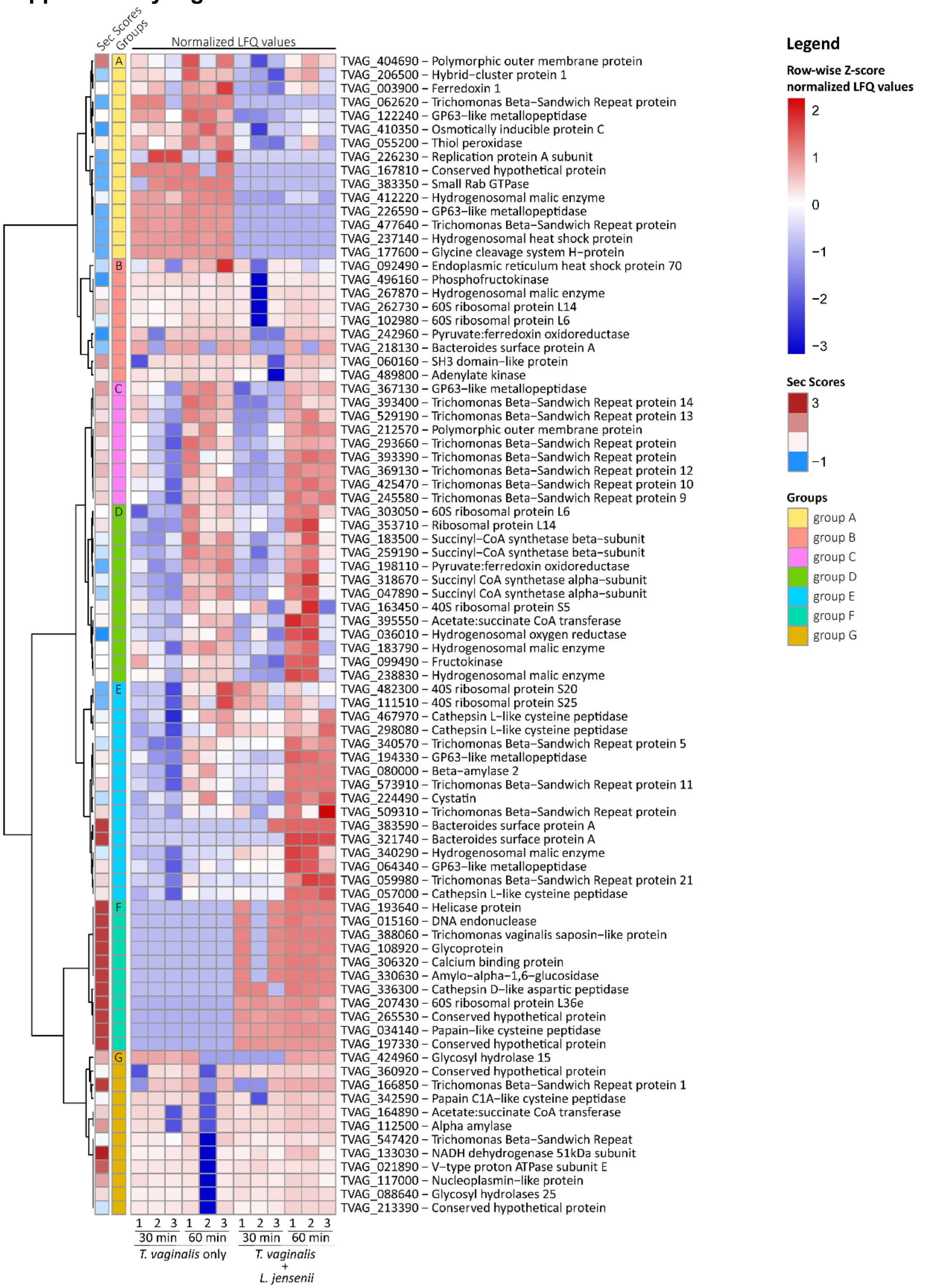


**Fig. S1.** Secreted proteins with a time-dependent increase. Seven groups (A to G) were assigned showing differences in secretion dynamics. Sec Scores, Difference of Secretion Scores.

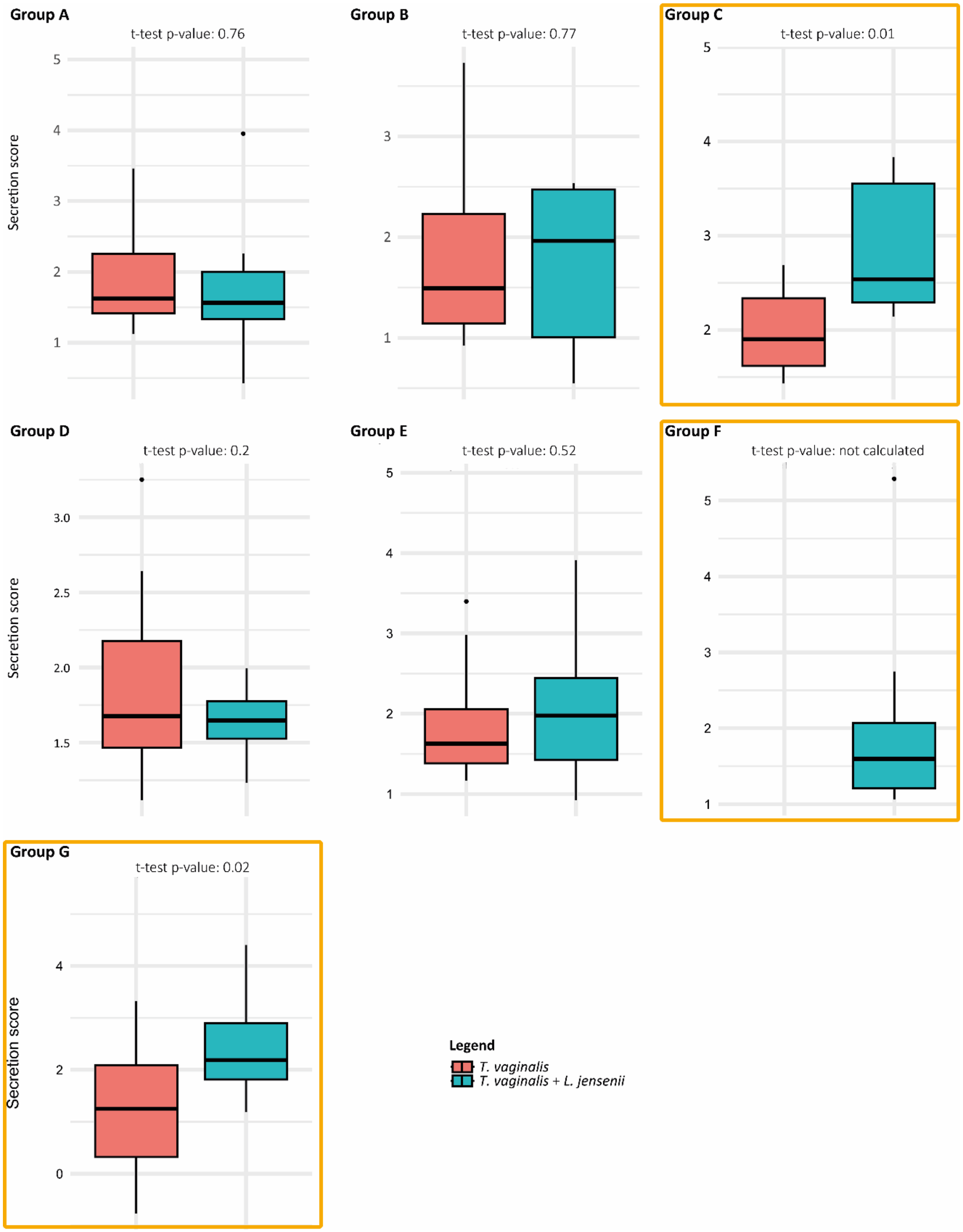


**Fig. S2.** Comparison of secretion scores between the co-incubation and control per group. Significance was tested using t-test, with the groups C, F, and G having a significant increase. Proteins in group F were unique to the co-incubation.

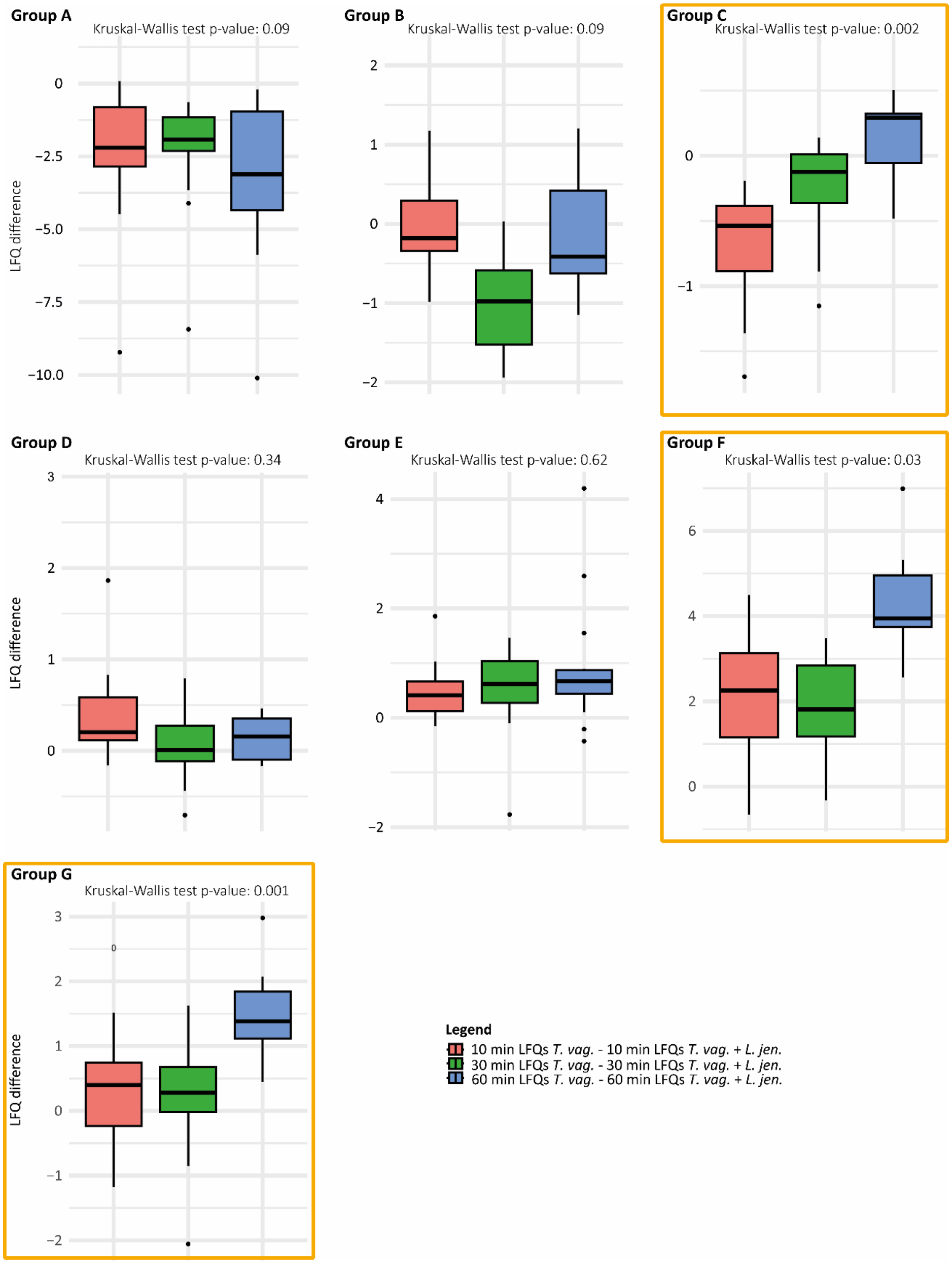


**Fig. S3.** Comparison of LFQ values between the co-incubation and control per time point and group. Significance was tested using the Kruskal-Wallis test, with the groups C, F, and G having a significant increase.

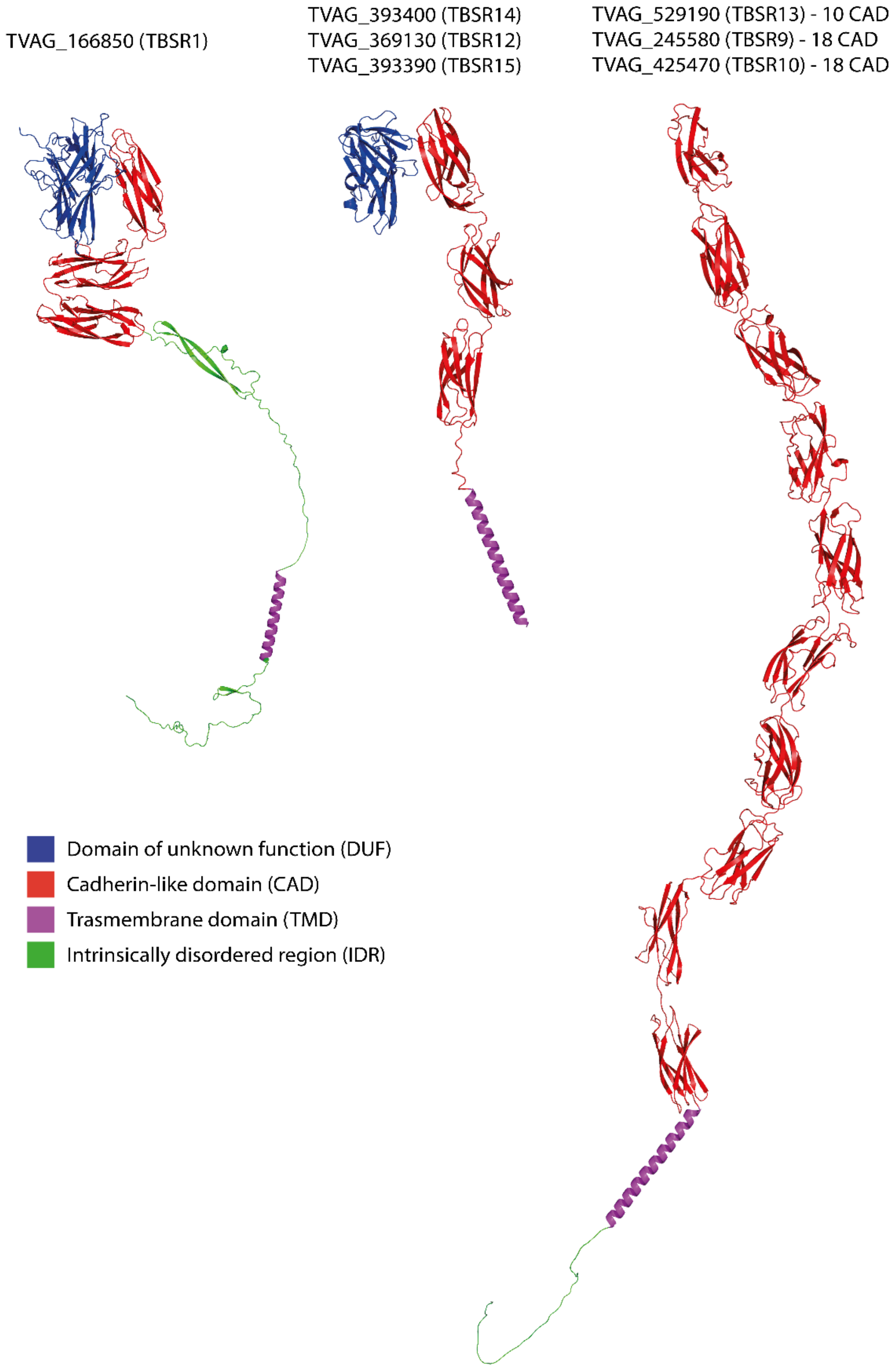


**Fig. S4.** Three-dimensional structures of TBSRs created using AlphaFold DB (8, 9). Domains are highlighted in colour based on The Encyclopedia of Domains (10).

```
TVAG_088640      ---------- ---------- --------MP YKVIDISVWQ GDIDFNAARD QSGVQGVMIR CGYGSYSPAQ  42
TVAG_122290      ---------- ---------- -----MVATP FKVIDISVWQ GDIDFNAARD QSGIQGVMIR CGYGSADPSQ  45
TVAG_368720      ---------- ---------- -----MPDLP FKIIDVSKWQ NEIDFQAAKD NSQIQGVMIR SSYGSYDPKQ  45
TVAG_015320      ---------- ---------- -------MNA YKIIDVSVWQ GNIDFNAAKV NSGIQGVMIR AGFGS-AAGQ  42
S.coelicolor     ---------- ---------- ------DTSG VQGIDVSHWQ GSINWSSSVK- SAGMSFAYIK ATEGT---NY  40
S.alcalophilus   ---------- MKLLPSLIGL ASLASLAVAR IPGFDISGWQ PTTDFARAY- ANGDRFVYIK ATEGT---TF  56
A.fumigans       --MKFSIVAI ATIAGLASAL PSQPEARATT VQGFDISNHQ KSVNFEAAK- KDGAQFVMIK ATEGT---TY  64
Caudoviricetes   ---------- ---------- --------MQ LSVIDVSEHQ GNIDWEKVK- NAGVVGAMIR TGYGVKDPNQ  41

                    α3          β4                    α4                β5        α5    α6
TVAG_088640      IDGQFHNHMR AAIAANMLIG VYHYGYAQSV PEAEGEADFA ISIFDQYRQW IRLPIAYDIE -DPTMNVGRD  111
TVAG_122290      VDKQFVNHIR AAIACNMLIG IYHYGYAENV QEAENEADFA LSIIEPYKQY ITLPVAYDIE -DPTMNVGRD  114
TVAG_368720      IDTMFENHMR NAIASNMLIG TYHYGYARSV EQAEREADFA ISVISPYKDY IKFPVAYGVE -DAAMNVGKE  114
TVAG_015320      EDKQFANHMR EAIAANILIG AYHYGYARTE AQAIAEADFC AKIINPYKSH FRMPVAYDDE -DATMNVGKV  111
S.coelicolor     KDDRFSANYT NAYNAGIIRG AYHFARPNAS SGTAQADYFA SNGGGWSRDN RTLPGVLDIE HNPSGAMCYG  110
S.alcalophilus   KSSAFSRQYT GATQNGFIRG AYHFAQPAAS SGAAQARYFA SNGGGWSKDG ITLPGALDIE YNPNGATCYG  126
A.fumigans       KDTVFNSHYT GATKAGLLRG GYHFARPDKS TGSTQAKFFL KNGGGWSDDN RTLPGMLDIE YNPYGATCYG  134
Caudoviricetes   IDKWFYNNLS GCQNVGMPYG FYHYSYAVDV AGAEQEADFC LELVA--ESN PQYPVAFDME ENRQAALGKT  109

                        α6                β6    α7     α8      β7
TVAG_088640      LVGQMAAA-- --FANKIRAA GYT-PMCYCN LNWATNFINM DIVNANGIDV WIAQYNSE-- ---CQYWGPY  171
TVAG_122290      LVGQMAVA-- --FANRIRAA GYR-AMTYCN LNWATNYINM DWINNAGIDV WMAQYNSQ-- ---CTYWGPY  174
TVAG_368720      LVADMVIA-- --FCNKVKAA GFK-PMLYLN LVWANNYCNM DRINEAGIDV WLAEYNSY-- ---CKYTGPK  174
TVAG_015320      QTAKNAIA-- --FCNRMKAY GYK-PMVYCN LNWARNYIDM SMINAAGIEV WIAQYNSV-- ---CQYTGPH  171
S.coelicolor     LSTTQMRTWI NDFHARYKAR TTRDVVIYTT ASWWNTCTGS WNGMAAKSPF WVAHWGVS-A PTVPSGFPTW  179
S.alcalophilus   LSQSAMVNWI EDFVTTYHGI TSRWPVIYTT TDWWTQCTGN SNRFANRCPL WIARYASS-V GTLPNGWGFY  195
A.fumigans       LSHSQMVAWI HDFVNEYHHA TSRWPMIYTT ADWWNRCTGN AKGFGDKCPL VLAAYSSSPP KTIPGDWKTW  204
Caudoviricetes   VCTDIAIA-- --FCNKIRAA GYQ-PMLYTN LNWATNYIDM ARIDAAGIDV WLAQYNTQ-- ---CDYKGSY  169

                     β8        β9
TVAG_088640      VMWQFTDCLP IGGYYAG-VD GSWCYKDY-- ----------  198
TVAG_122290      TMWQFTDAQP IGGYYAG-VD CSWCYVAYA- ----------  202
TVAG_368720      TMWQFTDALS ISGYPEP-VD ASWVYKNYE- ----------  202
TVAG_015320      TMWQYTNALK ISGYSAC-VD GSWVYKNY-- ----------  198
S.coelicolor     TFWQYSATGR VGGVSGD-VD RNKFNGSAAR LLALANNTA-  217
S.alcalophilus   TFWQYNDKYP QGG------D SNWFNGDASR LRALANGD--  227
A.fumigans       TIWQNSDKYK HGG------D SDKFNGPMTQ LRKLASG---  235
Caudoviricetes   TMWQYTSKGV VDGITANTVD MNWCYKDYTN GATTTPAP//  368
```

**Fig. S5.** Protein sequence alignment of TvGH25 with *T. vaginalis* paralogs and selected orthologs. *S. coelicolor*, *Streptomyces coelicolor* (cellosyl); *S.alcalophilus*, *Sodiomyces alcalophilus* (QNJ46261.1); *A.fumigans, Aspergillus fumigans* (XP_001481415.1), *Caudoviricetes, Caudoviricetes* sp. (DAY06394.1). Catalytic residues Asp[100], and Glu[102] are highlighted in red. Structural predictions are based on AlphaFold DB (8, 9).

**Supplementary Tables**

**Table S1.** List of sequences used for GH25 phylogeny.

| Species | Accession no. | Sequence |
|---|---|---|
| *Acanthamoeba castelanii* | ACA1_069730-t26 | MKSTSVLVVVAIVLASFAALPPAEAAKGLDVSLQDCASITPAQWRCLREQEGFSFAIIEAWNGGFQLNQKLAYCVSNAWAAGFAHVDIVHYYAFLCPNCGGNNPPANAVSAIDNYLKSNNVQYGQLWFDIEQCTGCWNDDASNFAFIKQAVASAQRLGMSVGIYSSDYEWGATVGASTRGFPGLPLWYAHYDNMPSFNDAWAYSFGGWTRPAIKQYYDRDSGACGVTNIDLDWYPDN |
| | XP_004347089.1 | MKSVLFALVLCSLIAASFATYGVDVSQETLPSAWSCLKQNGYSFGIVRAYLSIGQVDSVGPHTIYNAWAGGMSHVDVYMFPCPQCGKSAATQVQEAVQNLRNYNCQYGMFWFDIEGSQYWNDVASNRAWLTEAVNQALSMGQTVGIYTSQYQWSSIMGSWTGASRFPLWYAHYDNNPSFSDFSPFGGWNQPAIKQYVATPSCAALASTRTSTLERSRGS |
| *Acrasis kona* | KAL0484822.1 | MQNKLILLALSVFVVYANATLGLDLSFWQGDVSLNTWKCLRGSGYQFAIVQVWKQSGEPNPWAVSDIKRAYAAGFLHVDAYIFPKTSISPEDQVKRSIDLLRSQGAGNFGMIWIDVESKAAWGNCDQNVAYLNRLINQIKNMGVKPGIYSSSSQWSAIMCGRGGFSNIPLWYAHYDNNPSFSDFRSFGGWSKPAIKQYKGDTKQCGINIDANFY |
| *Acutalibacteraceae bacterium* | MEE0724001.1 | MEAKYKGIDVSKWQGDINWEKVRASGVEFAMLRAGFGRCQGQNDTSFERNYREAKKNGILVGAYHYSYAKTVSDAEKEAEYFLSLISGKSFEFPVAFDIEDNSQKNLSREEISNIVEAFCGKTEKAGYFVSVYANLWWLNNKISDRVKERYDIWLAQWADAPSYGGKYGMWQYTSSGKTDGITGNTDMDLAFKDYPNIMRANGLNGFSKGAAESTDNVKSGTFPPRRSVALCNTPLFSSAYSKAPSARKSGTYYIYDGIEINGRYRITSSASFALKKPIGKNVTGFVNADDIR |
| *Acytostelium subglobosum* | XP_012753434.1 | YIHTNMYTKLSAVILALVLIQVSVVFSSKGVDISVSSTIDDFKCLKGDGYSWVVIRCYESLGRVDSNCPHSIYNAKDGGITDVDAYIFPCTSCGNGAGQVVAMVNYIKSYKATVGRVWFDIEGPGTYWSNSHADNQAFFNSMLEGAKQAGVSVGVYTSASQWEPIMGSWDGGKALPLWYAHYDNNPSFSDFTSFGGWSKPYAKQYVGSTTVCGLGVDLDYY |
| *Amphimedon queenslandica* | XP_003384181.1 | MKMMFSALVVICSLFLGIQSLPVNRTAAGTYGVDVSSLISESTFSCLKSNGFEFAIIRAYRSTCSSDPNAPATIRNARAGGIKYVDVYMFPSPRCSKSAAEQVEEMVNSLAESNYGQIWLDIEGSQYWLGSESANRQFFDELLSAAKKYKKVGVYASQYQWSSIMGSYTGGSSQPLWYAHYDDNPSFSDFEPFGGWSTPTIKQYAGDKTVCGADVDKDWY |
| *Anaeramoeba flamelloides* | KAJ3439686.1 | MKLKLLLIILLILKITLATRGFDFSYFQGTPSVTDLKCLKNDGFDFGFVQAQRSNGEYNPYIKQVFKNLREAGLKYVDVYIFPNVKEDARGQIYKTISKLKEDGVLTDNMVWLDIEKYEWPSSHTTNIKFIEEMIDEVENQYKGGKVAIYSSKYSWDAITGGTTKFKDYQLWYAHYDNNESFSDFEPFGGWTKPNIKQYKDTTSICGTSIDYDYYP |
| *Anaeramoeba ignava* | KAJ5073150.1 | MKILQIIFSLVLASYVFATYGLDMSIYQGDPALSDFQCLKKDGYDFAILQAQTSNGGYNPYVKSQFEKCHQAEINYVDVYIFPDVSKDAASQIHTTVSKAIGDGVLTGKNMIWLDIEPYKWSSNHQTNIDFIAAMISECEKVYNIKPGIYSNWNSWSEITGGTTRFSSYQIWYPHYDGSASFSDFEPFGGWSKPNIKQFQGTTSICGTEIDKDFY |

| | | |
|---|---|---|
| | KAJ5071007.1 | MKLLFLIFGLLILQTVFSTSGLDISQADCGSISLSEWKCFAQNGKSFAIVEAFDGGYGIGSHVKDCVANAWSAGFAHVDVYAFMCPNCAGNNPPSSAVEKLVGYLSANRVNYGMLWFDIEQCSGCWNDASSNVAFIKEAVEKAQQLGVNVGIYSSEGEWSATVGSSSAFSNLPLWYAHYDGEPNFNDQWAYSFGGWSSPAMKQYYDSSSVCYSNVDVDWYP |
| *Ancylostoma caninum* | RCN53780.1 | MNEPDTIKMRKRCSFTHSIMRVATVLAALFGTSLAATGFDAIGAISTSHFSCMKSHGHSFFIGRVFRSSGTVDSQGIQNIKNAKAAGIAHVDGYIFPCTTSSCPSPATQISETSKALKSAGASVGMLWLDIERYQWPSDHTKNRQFIEAMGKELTSLGYKWGVYSNYNNWESIVGLDWTGMSSKQLWWATYNGQANFNNFKAFGGWSKPNIHQYNGDVTGPCSFSIDTNWYP |
| | RCN34231.1 | MKTLALLCSLALSCFASPVVQQQAASESASIAYAVDLSVPVSTSTFQCIRQSYYNVAFVRGYSPAGQGQFDPYACTNIQNANAAGLGTEVYMTPQPKSSKSGAQQFEEMYGGLKNSNINVRSVWIQVTSPINWYASSTTNINILNSILSRASQYGLAIGIYTSVYDWNQITGGATINNAMLWYWSVYGAGPTNESPANFNDFRSFGGWTAPSVKQFGQVESVCGVTVNRDIYTVSGAAKLAGMAKYEKSEQIVVGSLGLGTVAPGKAEIKQ |
| *Atopobium minutum* | WP_074667968.1 | MAMKGIDVSTWQGSIDFNRVKQSGINFVIIRAGYGSALSQKDKWFETNYARAKAAGLHVGAYWYSYAGSAGEAREEARICKQVLSGKQFDYPIYFDLEEKSQLARGRAFCDSLIRAFCNEMEAGGYFAGFYTSLSAALNYVSPDVRNRYAFWVAQWNSRCTYQGQYGLWQYSSSGSVPGIAGRCDMDLAYVDYPSIIRKGGFNGYGKGTSSAPARKSVDTLAHEVIAGAWGNGEDRLNRLTKTGYDYDAVQARVNELLGIKPKKSINALAREVIRGDWGNGKDRVNRLTKAGYDYEAVQKRVNELL |
| *Bifidobacteriaceae bacterium* | RFT36047.1 | MSKRGIDVSVWQGDIDFNAVKASGVEFVIIRAGYGIGHKDKWFEENYRKAKTAGLDVGAYWYSYASSAGEASEEAQSCVNILSGKSFEYPIYFDLEEKSQLNRGRAFCDSLITSFCSKLEACGYYAGFYTSLSVANNLVSSHVRDRYALWIAQWNTHCSYEGSYGLWQYSSSGSVSGIAGRVDMDYAYVDYPSVIKNAGLNGYKNGGSYTAPQTSSIDEVAREVINGDWGNGAERKQRLTSAGYDYASVQNRVNELLGVKAYRKSVDELAREVIRGVWGNGAERKQRLSQAGYDYDTVQKRVNELL |
| *Blepharisma stoltei* | CAG9314774.1 | MKSQALIILALTVIAVNATLGTDVSTAVSASQWSCLKSNSRVFTIIRGYQSFGQVDPNVVNNIKNARAGGQADVDVYIFPCVTCGNPAGQATALVDAIRGQNYGMVWIDIEVYQWSSSLTSNQAFITTMANTLKNLGQHVGIYTSYYNWQSIVGLSWSGVSWAPLWYAHYDNNPSFSDFQAFGGWSSPAMKQYNGSSTICGVGVDLDWY |
| *Bodo saltans* | CUG87019.1 | MTNSALLVAAVFIAVCAMTSNAFDGGDISEPYDASTFACCKNNGWGFVIVRSYRSYGAVDVNAPGSLDAAAAGGIEYRDVYHFPCTSVAASTQVSDDNAHVAGKYGMMWFDIETNYSPSCGWSGDKATNCNFMGALIDAGRAIGVHMGIYSSAYMWESIMGGCTVGADNGIPLWYAHYDYTRSFGDFSPFGGWTKPNAKQYNDAVGICGIDADADWYPN |
| *Cafeteria roenbergensis* | KAA0166630.1 | MRTQSILLALAAAALVPSALATYGADVSSHVSPDQWKCLKGDGINFAIVRAWYSFGAFDPNGRTTVANAWAGGMDDVDVYLFPCAGKSAADQVAGTVSGMSGVKFGQIWMDIETNPSPGCGWGSDLARNCEYIGELLQAAEAHGIKPGVYASPYMWGKIAGSACTVGRDYPLWWADYNGVPGWPPFGAFGGWSKPAMHQFADNGAACGVSYDKNYY |

| | | |
|---|---|---|
| *Carpediemonas membranifera* | KAG9391127.1 | MKLTFVVLLSLCVAALATKGFDFSVWQGTPSVADLECLKHNGYEFGIIQAQCSNGVYNSHV<br>AQAFRNAKAAGIKYVDLYFFPSKSHDAASQVRNTVNHLKADGLHSSSTQMWIDIENTGLFF<br>PSCSENVRFIREIVDNLRKEWPGKIGIYTSESQWSPITCNSKDFADMQLWWPRYDGVGTLT<br>HNWRSFGGWSRPAIKQYHNTMSICGTQIDEDCY |
| *Caudoviricetes sp.* | DAY06394.1 | MQLSVIDVSEHQGNIDWEKVKNAGVVGAMIRTGYGVKDPNQIDKWFYNNLSGCQNVGMP<br>YGFYHYSYAVDVAGAEQEADFCLELVAESNPQYPVAFDMEENRQAALGKTVCTDIAIAFCN<br>KIRAAGYQPMLYTNLNWATNYIDMARIDAAGIDVWLAQYNTQCDYKGSYTMWQYTSKGVV<br>DGITANTVDMNWCYKDYTNGATTTPAPQPEPSYDTYTVMAGDTLSGIAAKFGTTYQELAAI<br>NGIADPNVIHVGQIIKLKGDAASTPQGENTYTVQAGDTLSGIAAKYGTTYQELAALNGISDPN<br>VIHVGQVLKVTGSAGGRTYTVQSGDNLWDIAQSQLGDGARFREIIAMNGLLSDTIYPGQVL<br>QLP |
| *Cavenderia fasciculata* | XP_004357314.1 | MKFHALIFFIVSFLLLANVSLATKGVDVSQSISSSAFKCLKNNGFDFAIVRCGRSSGSPDPNC<br>ASNVQNAWGAGMAHVDAYIFPCPSCGNGASQVINTVRAYGGKSVGMLWLDIEGPQYWSS<br>NQGTNRAFFNSMLDGARQAGVRVGVYTSSSQWNPIMGSWSGGSAYPLWYAHYDNSPSF<br>SDFRSFGGWSKPAMKQYQGDQSQCGVGIDLNVY |
| *Circinella umbellata* | KAI7859614.1 | MKPSSPFTILASVATVLSFFSNIGVEAATGVDASSLITTEQWACAKETGYENAIIRGYIEAWG<br>QNPGGAVDKNVVENYKNAIAGGYTSVDLYIFPCTGRDTCKSPADQVQDVINLIDENTMDIGT<br>LWLDVEVDPEADNFPSAEGARTALQEFKKALDNSGEKWGVYASKVQWTSITGSVDWELD<br>SSVPLWYPHYDDVMSFDDFESFGGWTTPTIKQYAGTSNFCDAGWDKNFYA |
| *Clostridium sp.* | MBS5343358.1 | MKLSVIDVSEHQGNIDWKKVRSAGVSGAMIRTGYGVSSPNQVDKQFYNNLSGCKIVGMPY<br>GFYHYSYAMDVAGTEQEADFCLELIAGTAPQYPVAFDMEENTQAALGKAVCTDMAIAFCNK<br>IRAAGYTPMLYTNLNWATNYIDMTRIDAAGIDVWMAQYNIQCDYKGAHTMWQYSSKAVIDG<br>ITANTADMNWCYKDYTNGSAFIPTPKPKPSYDTYTVMAGDTLSDIAAKFGTTYQELAAINGIE<br>DPNMIHVGQIIKLKGHEISTPQGGTTYTVQAGDTLSGIAAKYGTTYQELAVLNGISDPNIIHVG<br>QVLLVSQNSSSHTYYVQAGDSLWGIAQSQLGDGTRYQEIKALNGLSDDTIYPGQALQLP |
| *Coriobacteriia bacterium* | MDO5334607.1 | MTQRSTFAQPNGIQQANAWPCALACALMAAFLFVALMFAAPQLAFADEDFEDETPTAEEV<br>EVPVESSEETDCANSWRFTNGEPTQESSLYRTTRASRAPSSKGSLGKLIDVSKWDGKINW<br>EKVRKSGIRYAIIRCGYGENKTSQDDKQFRNNVRNARAAGVKIGVYIYSYAYNTKKAQSEAE<br>HVLRLLKSEGLTPSKLALPVYYDLEEQAGTSKPCVKSDGKKYKVSNAMLGKMATTFAKTITK<br>AGYKVGVYANKNWWNNFLTDKAFSNKSWSKWVAQYSTSCTYKGSYDIWQYSSTGHVSGI<br>STYVDMNYLYRNFDTKIATASIGMTGITCPYQVVRGTNDLSLGGKLTSPVTLKTVTVRILEGD<br>SDAGAVKKSVTLYPGATTFDLASAGKLVEDLSFDGLPLGTYTYQVRAGVEDGQTTVFSKSF<br>KVVEPRSDGKTFLLATSSNVESAYLDYTRPTFGTSGLTLSTGLTKGGTLPLKGGKVTSSVAL<br>TRVQATLTNLTTGKQVAQWSGKASAKSLALPAACISKLSTKSLPMGKYRLVVTASSSSATG<br>TALSKTFTIGTAAPGVGKAKAAKKKFTAKWKKSSGATGYQVRYSRYKSMKSDYFKTTKSTS<br>KTIKVKKKKTRYYYQVRSYKVIKGAKCYGSWSSVKSVKVK |

| | | |
|---|---|---|
| *Corticium candelabrum* | XP_062513430.1 | MVMNLSVFIFCAAAYCLCGTVTARTLDQARAAPGGEGEKNQYKLPLFVRFQTGIFGVDVST<br>KTSTEAFSCLRRDGKEFAIVRAFRSIGKTDVNAPSTIASAWAGGFGRVDVYIFPCFSCGNPS<br>SQVTRTVDWLRNSRYGMIWIDVEQKKYWSDNQAANRKFFTEMLQTAQRYKPTGVYTSMN<br>YWTQIMGFDFTGAAHVPLWYPHFETAPSFSDFTSFGGWARPNIKQFDADNNVCGAVVDVN<br>WHP |
| *Dichotomocladium elegans* | KAI9315954.1 | MFAKTLLLGALTLLSATATQAYETGLDVSALTSTSSWSCARNLGYDYPIIRCWFEAYGQNPG<br>GALDKNCLQNYKNALAAGYSSVDVYMFPCSGRSTCKSPDTQVKELSSFIDSNGMKVGRIW<br>LDVEIDPAANNWPSPADAKTTLKAYKTALDASGKSWGVYASKSQWSSIIGDLSWELDASVP<br>LWYPHYDDVLSFDDFSSFGGWTKPTIKQYSGSVTFCSAGWDRNYYG |
| *Dictyostelium discoideum* | DDB_G0274181_t1 | MRFIALLISFFALLKVISAISGVDISSASTIESFTCLKSAGYDFAIIRAYESLGQVDPNGPHSVY<br>NARDAGIEYVDVYMFPCPTCGNGAGQAETMVNYLKGYNANYGMVWLDIEGPQYWMSQS<br>ENVAFFESLVAGLKAEGAHIGVYTSASQWEPIMGGYTGGSEFPLWYAHYDGNPSFSDFSP<br>FNGWSTPSVKQYDDTGDSCGLGFDLNWYP |
| | DDB_G0276439_t1 | MNKLILSILSVLLIVSIASAGNGIDISSGTNPSSSDLACLAKQNSKIIVQTWSGGYGFNSNVPSI<br>VQAAKSAGLSVDLYAFLCSQCSGITDPAEAIKTLVSKLNGVKYGTLWIDVEQCNGCWSSTL<br>SSNAQYVQTAVQTASQLGVNVGVYSSEGEWPQTVGSLTSLSSFPLWYAHYDNNPSFSDQ<br>GYNFGGWSKAVMKQYQGTTTQCGVSVDLNWFPGSGSSTTSSSATSSSTTGRPLTSSSGA<br>GTGGHSSSSGAGTGGHSSTTGGQTSGQTSGQTSGSSNQPLSSGSSQDSSNSGSSNQPL<br>SSGSSQDSSNSGSSNQPLSSGSSQTSGQTSGSSNQPLSSGSSQDSGSNSGSQGTTGSQ<br>SGSSSSNPLTSGSQSGSSGSQSTSSGSQSGSSGATGTSSTASSTSATGSSSTGSSTGSSS<br>GSNDSGSNDSGSSSGSNNSGSSSGSNNSGSSSGSASSSGSSGSSGSGNYTSGSGNGAF<br>LF |
| | DDB_G0293492_t1 | MRLFLLLITFIALFGAINAFSGVDISQGSSVGDFQCMLNQGFEFAIIRGYMETGQVDPEVVNSI<br>ACAREAGVEYVDTYLFPCFNCGNPQDQGPALVNYLSGYNANYGMVWLDIESSDWSGDQS<br>ANVAFFEGLISGLQSVGAHIGVYTSASQWIPIMGGYTGGSEFPLWYANWDGVQSFDDFSAF<br>GGWSTPAIKQYNDGGSNCGVGYDFNWYPN |
| *Dictyostelium purpureum* | XP_003289340.1 | MRLFLLLISLIFLAVQRTVAFSGVDISSGSDVPFFQCVKNAGFDFAIVRAYESIGQVDPNGPH<br>SVYNARDAGIEYVDVYLFPCFSCGNGAGQAVSMVNYLKSYNANYGMVWLDIEGPGTYWS<br>SNQGANQAFFNELVSGLESVGAHIGIYTSESQWEPIMGDWSGGAKYPLWYAHYDGNPSFS<br>DFSPFAGWSSPAIKQYNDQGLNCGVGADLNWYP |
| *Didymodactylos carnosus* | CAF0927709.1 | MTGKMIGLICFLTVSITTILATKGIDVSQSVTQAQFECLKNAGYEFVITRVHQSTGKVDPNGA<br>RTIKNAKAAGFRYVDGYIFPCFSCGNPAKQVSDTIDNLNSNGAEYGMIWLDIESGANWSSIA<br>QDNIDFIQQMIDELKRLKVNFGIYAQKSQWTHITGNTVVFGDPPLWYPHYDNVESFSGFQA<br>FGGWTKPAIKQYQGDVSECGVGIDRNFY |
| *Dientamoeba fragilis* | k64_S216140.p1 | MQSISAGKYKGIDISSWQGEVNFNQVSNSGVQVVYIKATEGDYYLNPRADEYYSGARSNN<br>MLIGFYHFFRPRVDAKTQANYFSSYVKGKSFNCRLALDIELSDGYGAAEISNMCVTFLEEVK<br>RLTGKDVVVYTYTNFARTSLTNILSKYPLWIANYDVTTPGDNPIWNSWVGFQYSESGTVPGI |

| | | |
|---|---|---|
| | | SGYCPLDVFTDGILISGGNTSIPNINQGGTTTTYTVKEGDTLSEIAQRYGTTTERLAQINNISN PNLIYPGQVLKISS |
| *Dysidea avara* | XP_065895301.1 | MLLTACVVLLLSLFGVVHVVEATKGIDVSTLVSSSDFQCLKKEGYDFLIVRAYRNIGKPDSNA VQTIKNARHAGFGHVDVYMFPCPQCSKPASVQVQEMVSALRGVSYGEIWLDIEKSSYWTG GYFGNQNFFEELLSEASKHKRTGVYTSKSQWNEIMGSSYTGGSSHRLWYAHYDNHMSFS DFTPFGGWHTPAMKQYAGDQTVCGVGVDLNMY |
| *Entamoeba histolytica* | EHI7A_001360-t26_1 | MFVFALLLALASAESYGVDVSQPTSASSISCLRNNGFTGFFICRAWCSPGYFDDNAPYTLG QAQSVGFVGDNSEVYFYPCLSCGNAAGQVQSFWSSVIANQMKFKRVWFDIEGQWYSDVG SNRAFFQELIDTALGMGIVSGVYSSQYYWGCIFGSGYSYGPAASLPLWYAHYDNWASFGD FSSFGGWSWPTMKQYRGDVSICSAGVDYNYRE |
| | EHI7A_061640-t26_1 | MKPDTFTCIKNKGYGDFVIIRGWRSIGSFDSEAANSYKSAITAGITDIDLYFYPCVKCGNAAS QVNSFITQANTNGIKWNTLWFDVEKDSWSSNVATNRQFLEDLLSAARAAGINHGIYSGWGV WGQVLGSSYVSKEASKLPLWFSHYDHKANFDDYNSGKYKFGDFTIAMKQYEGTTSMCGV LVDIDYKPAGVTPVNPTPEPETPTDGSTDGNSPSTPSTDGNTPSGDSTPSTDGNTPSGDST PSTDGNTPSTPSTDGNTPSGDSTPSTDITPSTPSSGSNTPSTPSSGSNTPSSSSNTPSSSS NTPSSSSNTPSTPSSGSNNGSSNGSNNTSKPRKTKEEKRLSKRIRKLKRQLYDKNGKLKEI KRLPGESFLHFLQRKAKQQKKIERLEKMEKRRDYLRKVRKEQRKINRMEKKLFDSNGKLKEI TQQPGESKESFLRRKRKQQRLLQRYEKEKKKENLLKWNNYKKSKHPLAELKGKLLDKSGK VKKLVKKTGESDAQFAKRVSKQKKLLKEYKKELKNSNKDKKDKKIWDKLKPKHKKKNGMF GKIERDINKSIKKTKKSIKKDIKKSKKQIKKIIKRKANKEGLNKKKTKKITKKQNKKMDKIKKGS NNRLKQLHKKSKKQLKQLEKKTKIKYQQAKRQGIQKAKQIKRNATRKALTIKRNTQKKAAAIR RMEKRKATIKKQQTKKTIIQRRNTKTRKSVKKVTVTRRR |
| | EHI7A_067550-t26_1 | MLFILAAIVLTAHAASYGVDIAESISTASIKCLRDLGFNSFFIFRAWRSLGSFDPNAKSINENAL AAGFSAKNIDAYFYPCYSCGDPAGQVKSFWNQVTSKKLKFERLWFDIEGTWSSSKSSNRK FFESLTNEAAAIGIKSGIYASYYNWEDIVGLDYKFSRAGSWPIWYPHYDGKKTFSDFKSFGG WTSPYMKQYNGDMTACSHDVDYNYRP |
| | EHI8A_022330-t26_1 | MFALFLCIALASAKLGIDVSQPTSTSSFTCLRNKGFTTMVIVRAWKSTGSFDTNAPQTLKNA NAAGFSIENSDVYYYPCISCGNMAGQVRTFWQKVGQYSLKVKRVWFDIEGTWTSSVSTNQ NYLMQMMNEARAIGIVHGIYGSKYYWGNLFGSSYKYAYASSTPLWYPHYDNSPSFSDFSS FGGWTSPSMKQYRGDVSVCSAGVDYNYKP |
| | EHI_081740A | MQVFFLCALIFTAFAGYGVDIAEPISVSSINCLRNIGFNSFFIFRCWRSLGSWDPNAKGINNN ALSAGFDAHNIDAYFYPCISCGDPAGQVNSFWNQVVANDMKFERVWFDVEGYWNDEESN KEFFETMINQQRAIGFKAGIYTNYNNWDQIFGLDYTFKYADEYPLWYAHYDSWDSFGDFTP FGGWSRPTMKQYNGDMTACSHDVDYNYKP |
| *Entamoeba invadens* | EDI_092550A | MFVFALLLALASAASYGVDVSQPTSASSISCLRNNGFTGFFICRAWCSPGYFDDNAPYTLG QAQSVGFVGDNSEVYFYPCLSCGNAAGQVQSFWSSVIANRMNFKRVWFDIEGQWYSDVG |

| | | |
|---|---|---|
| | | SNRAFFQELIDTALGMGIVSGVYSSQYYWGCIFGSGYSYGPAASLPLWYAHYDNWASFGD<br>FSSFGGWSWPTMKQYRGDVSICSAGVDYNYRE |
| | EDI_206270A | MQVLFLSILIFTAFAGYGVDIAEPISVSSINCLRNIGFNSFFIFRCWRSLGSWDPNAKGINNNA<br>LSAGFDAHNIDAYFYPCISCGDPAGQVNSFWNQVVANDMKFERVWFDVEGYWNDEESNK<br>QFFETMINQQRAIGFKAGIYTNYNNWDQIFGLDYTFEYADEYPLWYAHYDSWDSFGDFTPF<br>GGWSRPTMKQYNGDMTTCSHDVDYNYKP |
| | EDI_273010A | MLFFTLLIASVFAIRGVDMSTLTSVSSFQCFVNNGYTFAIPRCWRSTGSWDPYCANNCANA<br>HAGGMSRVDAYFFPCYSCGNAAGQVNSFWQKVVSNQLKITTVWFDIEGTWSSSYTANQQ<br>FFMEMMNQARAIGIVHGVYVSQHYWNSFFGSSYIYPYASSTPLWYPHYDHNPSFSDFSSF<br>GGWSTPAIKQYQGTTSMCSASVDLSYQP |
| | EDI_281570A | MLFIFAAIILTAHAASYGVDIAESISTSSIQCLRNLGFNSFFIFRAWRSLGSFDPNAKSINENAL<br>AAGFSAKNIDAYFYPCYSCGDPAGQVKSFWNQVTSKKLKFERLWFDIEGKWSSSKSSNRK<br>FFESLTNEAAAIGIKSGIYASYYNWEDIVGLDYKFSRAGSWPIWYPHYDGKKTFSDFKSFGG<br>WTSPYMKQYNGDMTACSHDVDYNYRP |
| | EIN_086570-t26_1 | MFFVLLFISAALAGKGIDMSTAFPTSTFSCFVSNGYTHAIPRCWRSSGSWDPNCASNCKNA<br>HSAGMSQVDVYFFPCYSCGNVAGQVSSFWNQIISNKVDCKRVWFDIEGTWSSSASTNQN<br>FFMEMMNEARAIGMVHGVYVSKYYWGQFFGSSYTYKYASSTPLWYPHYDSNPSFSDFAA<br>FGGWSSPLMKQYQGTTSMCSGSVDFNYF |
| | EIN_284410-t26_1 | MLLFACLLSLSTALLGTDTTYVLPTSGIACLKNMSSDFIIFLASDSNTGKFYSSVASTVKNAW<br>SIGFEKAQVDLLFTPCVTCGNPETQIVEFYGNVTSNKLKPNCIWLNVIDGWYSDIPDRQGFF<br>EKMIAQATKSKMTFGIFTSKKYWEGVFGDTYECPNARTTPLFYYNSDNNPSFSDFRRFGG<br>WAKPTAKRYSPNNYVCNTKANYDYKE |
| *Entamoeba moshkovskii* | EMO_019210-1 | MLVFALLVALVSAESYGVDVSQPTSASSIQCLRNNGFTGFFICRAWCSPGYFDDNAPYTLG<br>QAQSVGFSGENSEVYFYPCLSCGNPAGQVSSFWNSVVANRMNFKRVWFDIEGQWYGDV<br>GSNRAFFEQLISSALNMGIVSGVYSSNYYWGSIFGSGYSYGPASNLPLWYAHYDNWASFG<br>DFSSFGGWSWPTMKQYRGDVSICSAGVDYNYRE |
| | EMO_020450-1 | MLVAILLTLFIANANAARGIDCAYSMTVDTFKCLKEKGYGDFVIIRGWRSNGKFDTNAPKSYA<br>NAKAAGITDIDIYFYPCISCGNPSSQLNSFLSQAKANGISYGRLWIDVEVMSWSKNQQTNRQ<br>FLEDLMAACRAAGVTHGIYSSAGVWGPVMGQGYVSKEAAKLPLWYSHYDGVESFSDFSK<br>EKYRFNTYTSPSIKQYKGDQKLCGAGVDYNWKPGASAGPTKEDEDKKKKEEEAKKAEEEK<br>KKKEEEAKKADEQKKKVDEEQKKKQEEEKKKKEEEAKKADEQKKKADEEKKKADEEAKKK<br>PQQEKTEKEKIKEEAEKEKQKKVEKERKKEEQKTKDERALKEKEKKQKENIKRKVEENQNE<br>KKKEIKPKKDKTVGGFSSNAEKEAAAKGKMGGSSKKKNKLDSISDKTGSYVDKTTKRTKKE<br>LNKIIGRKAINDMKKQYEKTKELTQKGLKKVTKYIKKGKQVAKTTQKWLNKGKSKIDTIGKKA<br>KTFVKKQTKKLEHLKQNTLNKAEELKQRTSQKWKQLTQDTRREISNAKKISLQKVQEAKAIA<br>QRKALELRERTLEKARQLKLKAQEKARRYREAAIKKANDLKQKALAQKQKAQQKMKELQIQ<br>AQKRAQQAKKKMAETKAKLIKKGNELKNKAFTATKNTYASAKKKLSKIF |

| | | |
|---|---|---|
| *Gardnerella swidsinskii* | WP_116440732.1 | MSKKGIDVSVWQGDIDFNAVKASGVEFVIIRAGYGIGHKDKWFEENYRKAKTAGLDVGSYW<br>YSYASSAGEAALEAQSCVNILSGKSFEYPIYFDLEEKSQLNRGRDFCDSLITSFCNKLEACG<br>YYAGFYTSLSVANNLVSAHVRDRYALWIAQWNTHCSYQGSYGLWQYSSSGSVDGIAGRV<br>DMDYAYVDYPSVIKNAGLNGYQNGGSYTAPQTSSIDEVAREVINGDWGNGNERKNRLISA<br>GYDYASVQNKVNELLGVKAYRKSVDELAREVIRGTWGNGNTRKQRLTQAGYDYNAVQKR<br>VNELL |
| *Gardnerella vaginalis* | WP_065189629.1 | MSKKGIDVSVWQGDIDFNSVKASGVEFVIIRAGYGIGHKDKWFEENYRKAKTAGLDVGSYW<br>YSYASSAGEASEEAQSCVNILSGKSFEYPIYFDLEEKSQLNRGRDFCDSLITSFCNKLEACG<br>YYAGFYTSLSVANNLVSSHVRDRYALWIAQWNTHCSYQGSYGLWQYSSSGSVNGIAGRV<br>DMDYAYVDYPSVIKNAGLNGYQNGGSYTAPQTSSIDEVAREVINGDWGNGNDRKNRLISA<br>GYDYASVQNKVNELLGVKAYRKSVDELAREVIRGTWGNGSMRKHRLTQAGYDYDAVQKR<br>VNELL |
| *Halicephalobus sp.* | KAE9546249.1 | VSESGFKCLWDHSYRFFVARVYRSTGAYDTTGIQNIKNARAAGWKDVDGYIFPCLSSSCPS<br>AKTQVANTVNELHAKGAQFGMIWLDIEKLAWPADHNHNRAFISEMMQELDALKVNYGIYTN<br>NYMWGSIVGDWSGGSHRPLWWADYNGRQDFSGFKPFGGWSKPTIHQYSGSVNGPCGV<br>NMDQSWYP |
| *Halteria grandinella* | TNV75753.1 | MCATVALAMRGVDYSTLQSVETHQCWKNTGLHFAIPRAYKSFGAFDTNGPRNVANARAAG<br>FEYVDIYMFPCRGKDAASQVADLVNQMGNSNYGQIWLDIETNESTGCSWAGHSGDSNCA<br>YVNELISAIKSHGKVPGVYSSYYMWESIMGHPLNCGGHNGIPLWYAHYDNSQSFSDFHSF<br>GGWSKPNIKQYAGTTTLCGGGVDFSWYP |
| *Heterostelium album* | XP_020427338.1 | MGTACNIFSNIFFIGTRDTLSESNLIFWYRQIVVFALCFIALFQVSNAAIGVDISSGSDIPAFQC<br>LKNNGYTFAIIRCYMSTGATDPNCPHSIYNARDGGMEYVDAYIFPCFSCGNGAGQVVDTVN<br>YLKSYNADFGMVWFDIEGPGTYWSDNQGENQAFFNSMLEGASQAGVKVGIYTSYSQWEP<br>IMGNWDGGAKYPLWYAHYDGNPSFSDFSPFGGWSSPAIKQYNGDLTVCGLGVDQNYYP |
| *Hexamita inflata* | CAK8747467.1 | MLITFAITLQTRGFDLSYYQGDVSQDSMNCLFNNGYRFGIIEAQIGNTFNQNSISDYQRAKA<br>AGFQNVDFYIFPTINKDARSQVRDTVQRLLNAGVMTGNMIWLDIENIDLFYPTQAQNQQFIA<br>AMLSEMSNMLGANRVGVYSNWNQWSQIVGSSWTGAQPHQLWYPHYDNWQSFDDFQPF<br>GGWTRPSIKQYLGDNPECSTHVDRNFY |
| | CAK8747859.1 | MIITFIASIQTLGADISERLSNIHAKCLFENGYHQIVVQAQYGSSINQEAIKNYEVATAAGIKRV<br>DYYITPSTANDPVEQIRKTMDVLQSHNVIQTNSIVWLNAEDIDSYFGTQRENQVFILRLLEEIS<br>QHLGSERVGVYTSNQSWMALVDPNWRDTIIYKLKYAHFDGVTDFDDFVSFGGWKKPSMK<br>TFTGDVNECGMRINKEYQ |
| *Kipferlia bialata* | GIQ84655.1 | MCEYVIFEAQRSNGKYNKYIKDWYPEAKGVVMSRIDYYISIDQKIDAAQQIQATINSLHIDRAL<br>TANTFFWIDVQNGQHWGSNTAANVQFIEDMVNAGNTAMAGLFSGAGANFGIQTSKSMWK<br>RITGDNRDAFFGGLRLWYVDWTGNGDMSDFKRFGPWYQPSLKQYQGNVGMCFTHVNYD<br>SY |

| | | |
|---|---|---|
| *Lactobacillus delbrueckii* | WP_268886306.1 | MKMGIYKGIDVSLWQKNIDFNAIKASGIDFVIINAGYGHSIHQKDPYFEQNYSRAKAAGLKVGAYWYSYADSATNAKLEANTCLTAIAGKQFDYPIYFDLEEKSQFKQGSAFCSGLVRTFCSEIESYSYYAGLYISRSPLQNYISPDVASRYALWIAEYNPRLNYSGNYGIWQYSSTGSIAGISGNVDLDYSYIDYPTIIKQKGLNGYGTTHPLKSTDEIVAEVLAGKWGNGPDRQNRLSAAGYNYSSIQASVNQRLNQKSIDTLAREVIQGQWGNDPQRSQKLRAAGYDPTAVQRRVNQLL |
| *Lactobacillus gasseri* | WP_235077138.1 | MISGIDVSEWQGHVDFNAVKASGVKFVLIRAGYGRSASQEDRYFAEHYIQAKAAGLQVGAYWYSYAVSSADAANEARACLTVLGNRHFDFPIYFDLEEKWQFANGRNFCDSLVKSFCSVLEQNGCYAGLYISRSPLQNYISPSVAQRYAVWVAEYGPCCNYNGNYEIWQHSSTGSVPGVNGNCDLDYAYIDYAAVINKKQPVTRKNPDELAIEVLNGQWGNGTDRQQLLTAAGYDYAVVQEKVNRLLNRKSVDQIAREVIRGSWGNGNERINRLSKPAMTPPKFKNELINYCNFCLWTPVCRLFSFTMVYFSACSGLSVEVIKYGKESTTSNPSTTNNNK |
| *Lichtheimia ramosa* | CDS02619.1 | MKNLLVGLSLLASAFVSTQAANTGVDVSALTSTSAWSCAKNYGYSHAIIRCYFEAWGGNPGGALDSSCAQNYANAVAGGFNQIDLYMFPCTGRSTCKSPATQVNEMVAHMNKNKMKIGTLWLDVEVDPQSNNWPSASQAQSTLKQFKQAFDATGLKWGVYASQSQWTSITGSKDWVLDSSVPLWYAHYDEALNFNDFSPFGGWTKPTIKQYAGSQSFCSGNWDKNFYG |
| *Ligilactobacillus animalis* | WP_225362238.1 | MKMGIYKGIDVSLWQKNINFNAVKASRIDFVIINAGYGHSIHQKDPYFEQNYGRAKAAGLNVGAYWYSYADSATDAKLEADTCLTAIAGKQFDYPIYFDLEEKSQFKQGTDFCSELVKAFCSEIESHGYYAGLYISRSPLQNYISPDVASRYALWIAEYNPRLNYSGNCGIWQYSSNGSIAGISGNVDLDYSYIDYPAIIRKKGLNGYGANTKPLKSTDEIVAEVLAGKWGNGTDRKNRLSAAGYNYSSIQAAVNQRLNQKSIDTLAREVIQGKWGNDPQRSQKLRAAGYDPTTVQRRVNQLL |
| *Limosilactobacillus mucosae* | WP_180304701.1 | MGIYKGIDISLWQKNIDFNAVKASGIDFVIINAGYGHSISQKDPYFEQNYSRAKAAGLKVGAYWYSYADSANDAKLEADTCLTTIAGKQFDYPIYFDLEEKSQFKRGSAFCSGLVRAFCSEIESHGYYAGLYISRSPLQNYISPDVASRYALWIAEYNPRLNYSGNYGIWQYSSNGSIAGISGNVDLDYSYIDYPTIIKQKGLNGYGTTPTLKPTDEIVDEVLAGKWGNGTDRKNRLSAAGYSYSSIQASVNQRLNQKSIDTLAREVIQGKWGNDPQRSQKLRAAGYDPTAVQRRVNQLL |
| *Monosiga brevicollis* | XP_001745316.1 | MLRLVLLAACVAAAAAVGGVDVSQITLPSAWSCLKENGYTFGVIRCYESVGQPDSACPHTIYNAWDSGMSHVDVYMFPDPSAGNPQGQVSTMINYLAQYSIKSGGSPPGAFGMIWFDIEGTQYWSSNQQSNRDFFSGLVSEAKALGLSIGVYTSESQWSPIMGDWNGGAAFPLWYAHYDGNPSFSDFSPFGGWSRPAIKQYGGTRSICGASVDVNWYPDGSSVFANRTYAHPHHNARNQSLVH |
| *Naegleria lovaniensis* | XP_044551439.1 | MKFHVVFCAVAALVLLGFLNALMDSKVEATLGLDLSYFQGDVPQSSWNCLKQNGREFAIIQAQTQSGHVNPYIVNDINRAKAAGIKYVDIYIFPNVNMDPAEQVRNTVSFIKQNGGTFGQIWIDVEGPQYWSSSCNNNVAFISKMVNAGHAAGYPTHIYTSNSQWQPITCGSTAFSHLQLWYAHYDGRASFDDFYAFGKWL |
| *Ochromonadaceae sp.* | KAJ1388145.1 | MLFALIVIFAFACRIASATYGVDISQRTYESSFSCLVSNGYIFAVPRVYQSSGVPDPNGPYSIEDAWAGGMSYVDGYIFPCYSCGNPAGQMDDTISYLSAHNVKVLKKGELRPVASGNSTEKLG |

| | | |
|---|---|---|
| | | ATVGMLWLDIEGTQYWSTNTANNVNFLTSMANEGKARGYSIGIYTSASQWNPIMGGSTAFSSYPLWYAHYDNSPSFSDFSPFGGWSYPNIKQYAGTTSICSASVDKNYY |
| *Paramecium octaurelia* | CAD8159344.1 | MKQFNQILFLIVIVVPTVIAVKGVDLSASFNNFFCIKNAGYQFAIVRGYMSYGAADTVGLQNLRNAKASGLIGDAYFFPCKGKKTASAQVSEFDDVFTSEYGTVWIDVEYNPSTGCGWSTSDYSGNCQFLQDIVRELGNRGRLVGIYASQYQWSTIMGSASACAYFTSKPLWYAHYDNSPSFSDYSKYSFGGWSTPSMKQYQGDTTLCNVGVDLNFY |
| *Paramecium sonneborni* | CAD8087874.1 | MKMTSLIIVTLQIYSSLAVLGIDLSAKHDDFGCFRMNGYQFAITRAYRSYGEVDLDGYENLEKAKAAGLIGDVYFFPCKGKKTAKSQVQEFVQVFGNQQTQVYGTVWVDVESNPSTNCGWTNDYASNCNFLQEIVNELKGNSRSVGIYASQYMWTSIFGSTSACSHFTSLPLWYPHYDHSPNFDDFPKYKFGGWTTPSIKQYSGTTNICGVGVDLNYYM |
| *Paratrimastix pyriformis* | KAJ4456822.1 | MQKLLLAALFVGVAFATLGVDVSQNFYVPVFKCLHDNGYDFAIIRAYCSTGRPDDSSVHTVYNAWEGGMKHVDIYMFPCPKCGNPAGQVRDAIAHQRAYNTTYGMFWLDIEEPSYWMDQGSNRNFIQGLIDEARAEGQVVGIYTNLYQWSSIAGDWSGAAGYPLWWAYYDGSPSYDHWQNFGGWSKPAIKQFQGDATVCGADIDRNWYP |
| *Pelomyxa schiedti* | KAH3756411.1 | MKRRLVTGFLLVCLWVGISECVTGVDISLYAEVQPSDWECLYKNGFRFATVQCWNCGYYFGEDTAYAVQTAWDAGFEAVDVYVFMAPKCIHNSNATLVVDWIYGNLTAQAINYGKIWFDIEECPSSSCWQSTSANAQYMHEAAKRADSLGMTVGFYSSVYEWGVVMGSDASFSSYLQWYAHWDENPSFSDTPKYNNYGGWVTPYMKQYWDAGPCSGIDTDVNYRDNI |
| | KAH3744709.1 | MRLCVVGIVLVCLVGLEQVYCYSGIDMSSTLVEANAECLVSKGYHFTVSRCFRSNGEVDTAVVASSKAAWNGGMSYFDVYFFPCFSCGDPAGQVSTSVQYLTSNSVKYGMMWFDVEGPGTYWGNSTSANQNFFTSLVAQAKSMGVTIGVYTSKSQWQPIMGSSYTGGAAYPLWYAHYDNNPSFSDYVAFGGWTTPSVKQYEGTTSLCNQSVDLDWYP |
| | KAH3756076.1 | MVALTLVFVFSAFCLQLSLATSGFDHSYYQGAVNTSSYECLFSGGYRYAIMQAQTGHPPGTGHYNPYIGENYHNAVSAGIYPVDVYIFPDYGLDPAGQVNSTITSLFEDDVLYHDNMIWMDVEGTEYWIGTCDENIQFLHTVISTAEAMYSDCGLKYCIGIYSSASQWSPIMCDTTEFSSYPIWYAHYDNNPSFSDWKNFGGWTSPVMKQITGTTDLCGTQIDNDYRP |
| | KAH3765864.1 | MDVSSLITADDFSCGLADGYDTVVVRAYHSYGGVDTNAPPTLENAQQAGLLVTDVYHFPCYGKDKTGELPAVQQVSDMVEYLASTNSTYGRLWFDVETNPSDGCGWETSSANFPLNCQFLGDLITAARDLNQTVGVYVSSYEWSIIMGSACTIASDTLLWYPHWDDNLSFSDFVPFGGWTEPTMKQYSDKEHICSLNEDADWWP |
| *Pentatrichomonas hominis* | k48_S457728.p1 | VSVHQNDIDWQAAKDSGVNAVMIRSGYSTDGPDKKFLLHIRSALAVKMAVGIYHYSYAMSVEEAAQEGAYCVSLIREFKDSITFPVCYDIEESKIAALGKDTTTAMARAFLNEVKKEGYTPMVYCNLNWANNYINIPELQAEGIKFWIAQYNSECNYKGTYSMWQYSSSTHLPGSDQNIDSSHCYEDFTGGTIDFKVNTQRGISPKMEFTEIPDVDLADLQKECGVTQNKVAGEGFLKVLMQHEVGLGDSGFFVEWVQKRLTKLLFYKGKTVSHVDGPLQKAIDAFHAYYQLPEGKISG |
| | k64_S227825.p1 | STPYKVIDISAWQGDIDFKATKASGIRGVMIRSSYGTENPKQIDKQLANHATTAIAAGMLVGFYHYSYASNIQEAEAEADFCLKCIAPYKKDTVLPVAFDLEENSVAGLGKDYLAKMSIAFCRKIK |

| | | |
|---|---|---|
| | | AAGYKPMLYTNLNWARNYVNMAMIDAEGIDVWLAQYNSVCQYTGKHTMWQYTDHERIAG<br>YGSGVDCSWCYVNYNDPDSPQMEQTQRGPNPSLVFNGKLDFYLTDLQYIYNLKQTAKADS<br>TLYTKVKGETIKQGTSGQLTRWAQHRLSQLGFRCAESGTANSETMASIKQFQAYYQLGQG<br>ELGGTDWYYLLQP* |
| | k64_S356992.p1 | SFRILDVSEHQDTINWTDAKEAGVKGVIIRAGFGSRNPSQIDKKIHNHALGALSANLAIGFYH<br>YSYAKSVEIAKNETDFFFDIISEYKNRTTLPVAYDLEDKTIINWGLTKEDTATMAINFATKARS<br>LGYIPMVYCNLNWAKNYINMKRVHEAGIELWIAQYNTHPQYIEGPIGMWQYSSKTHLPSYT<br>EKFLDSSWCTRDYTNGTISFRENKQRGPNPKLEFRNNLDIDLGDIQSDLGVTKLRIANADSY<br>YALTQYSVSLGTESFVVPFVQKRLKLLKFYDGDITGKADSATIQAINQFQKYYGLGQGTLQG<br>TDWYYMIQP* |
| *Phascolomyces articulosus* | KAI9250945.1 | MKPFSPFTILASAATVLSLFSNTGVQAATGVDVSTLISEKTWACAKDYGYEHAIVRGYIEAW<br>GQNPGGALDKNLVQNYKNAIAAGYETVDIYIFPCTGRDTCKSATQQIQDVLDVIDENSMKIG<br>TLWLDVEVDPQADNWPTASGARSTLQAYKKALDNSGEKWGIYANKVQWTSITGSSTWELD<br>SSVPLWYPHYDNERTFDDFEAFGGWTKPTIKQYAGTSNFCSAGWDKNYYG |
| *Polysphondylium violaceum* | KAF2072606.1 | MKVILLLCILTLLKAANATNGVDVSSLTSTSSFGCLKSNGFNFAVVRCFQSVGRVDPNCASS<br>VNNAWSAGMSDVDLYMFPCSSCGNGGGQASKLISYLKENNVKYGTIWIDVEGPGTYWGS<br>STSANVAFFQDIVSGLEKEGVNIGVYTSASQWAPIMGDYEGASKFPLWYAHYDGVTSFSDF<br>SPFSGWSKPYMKQFMGDVSTCGTTVDKNWYP |
| *Pristionchus mayeri* | GMR56339.1 | FQAMNRIAVLFLALGVSLSTANLGIDTIAAISSSGFSCLKSNGYHFYIGRVGQSNGGIDKTGIQ<br>NIKNAWAGGMTNVDAYLFPCHSSTCKSAKAQVQDTVNALKNAGAKFGMLWLDIEIYNWGT<br>NQNTNRQFVLDMVSECNALGVKPGIYSNNNNWQSIVGIDWNGVSKYPLWWANYNGQANF<br>NNFKPFGGWSTPAIHQYSGDVKGACSVGNVDLNWDP |
| *Pseudoruminococcus massiliensis* | WP_332017960.1 | MKEKLIDVSTWNGNIDWNKVYKSGVKYAMIRSSFGVENPNQIDNKFVRNITNAVRAGVKCG<br>VYHYSYAQSVAEAQKEAEFCLKTIKGYKIDLPVAFDIEDSSQTGLGRRVCTDMVIAFCDTIKS<br>AGYKPILYCNPNWLQNYLYKDELLKKYDIWLANWGVYSPSYDCAIWQYSENGSVSGISGSV<br>DMNWIFKDYTNSSNNNNKSSAVKVINKTSYLRSHPWVDDKGGSSRKLATLTVGTEVDYISD<br>DGCGWSKVHGNNTTGYIQNTRISKTGISAYRYVNIKIPLYSNISDKKPVTYLTSNDKYYYICSI<br>ESGKNAGWSIINANGKDYYIKTENIAYGSKRMK |
| *Rhizomucor pusillus* | QNJ46208.1 | MKFALLVSAIAGLAATAVQAYETGVDVSALTSTSAWSCAKKLGYDHAIVRCYIEAYGGNPAR<br>GKIDSNCFQNYKNAKAGGFTSVDIYMFPCTGRSTCKSPAAQVKEVVDYVGSNKMTVGRLW<br>LDVEIDPSANNWPSASSARSTLKSFKSALDSTGWKYGIYSSASQWSQITGSSSWELDSSLP<br>LWYAHYDASLSFSDFSPFGGWTKPTIKQYAGSVSFCSAGWDKNYYG |
| *Rotaria sordida* | CAF0878169.1 | MFIQIILISIVPTILATLGVDVSQRVSQEQFACLKESGYKFVIVRVYRSVGVVDTNSAQTIKNAR<br>AAGLNNVDAYLLPCISCGDAPQQVIETIDYLRDEEAQIDRLWFDIEGQWNNDTDINIQFVDEL<br>IKQTSTLDVKFGIYTSRYQWISIMNDATKFSSNSPLWYAHYDNDQSFRDFQSFGGWTQPLIK<br>QFISDVKECGMILDKNFS |

| | | |
|---|---|---|
| *Ruminococcus sp.* | SCJ23317.1 | MDKRGIDVSSWQGVIDWDAVKSSGVEFAILRSSFGSPSPSQVDNQFYNNVKGAQAAGIPIGAYHYGYAVTEEEARNEAGFFLDTVKGIRFEYPLYYDVEDSATMGSLDRDALTKVIRAFCETVEKAGYYVGIYASLNWLTNKFYPDQLPYDVWVAQYYSEDQYDGHHGMWQYTSGGTVGGIAGKVDMNIAYRDFPKLIKDKGLNGWGSGESGGSGGSEEGSGGERDDRLGIGHCVADVLNVRSGPGTDYPVVYQIFTGNMVDVLKISENGWLQINCLHGVGWCAAQYIQWSPFESQQPPIGIGRCTADVLNVRTGSGLDYPAAFQLSQGNMVDVLTASGQWLQINCLLGSGWCASQYIDWFRTNLQVPAIGVGKCTADVLNIRSGPATDLSVLFTISEGNMVDVLEDNGRGWLRIRCLLGTGWCSAQYIDWSRHDK |
| *Salpingoeca rosetta* | XP_004988233.1 | MKVFAFAAVVCAALVAVALCPMGAEATYGVDVSQQTYENAWSTTCITRAALNFPHVHTPQVQRVRFGVIRCFMSTGHPDPNCPHTIYNAWDGGMDHVDIYFFPCPTCGNPAGQIDAMVANLAKYNIKPGNAKPGTYGMLWLDIEGPQYWTKDVGANRNFFSGLVSQAKAHGIHLGVYTRQSQWSPIMGDWSGGSAFPLWYAHYDGSASFSDFRAFGGWSKPSIKQFAGTTSLCGAGVDKNWYP |
| *Streptococcus anginosus* | MED5846031.1 | MTIKGIDVSTWQGSIDFKRVKQSGINFVIIRAGYGSALSQKDKWFETNYARAKAAGLHVGAYWYSYAGSAGEAREEARICKQVLSGKQFDYPIYFDLEEKSQLARGRAFCDSLIRAFCNEMEAGGYFAGFYTSLSAALNYVSPDVRNRYAFWVAQWNSRCTYGGQYGLWQYSSSGSVPGIAGRCDMDLAYVDYPSIIRKGGFNGYGKGTSSAPARKSVDTLAHEVIAGAWGNGEVRVNRLTKAGYDYDAVQARVNELLGIKPKKSIDTLAREVIRGDWGNGKDRVNRLTKAGYDYEAVQKRVNELL |
| *Symbiodinium sp.* | CAE7188006.1 | MKTALSGAVVLAAISSALAARGVDVSSHIQPSTWTCLAEEQYEYAIIRAHLGSGGPDPVAPHSIYNWYSSLMYKKGGRLGIYLFPNVPRGEPEAQIAETIEYLKGYSITYSAERGLHTFDRIWLDIEPQAWSSDKAANAEFIAGLVKGCEEAGVQCGIYANWNSWAEIAGEWSGASHLPLWYPDYDGTESCSGFRPFGGWGRADYKQWQGDDTICGAEVDLNTGC |
| *Tetrahymena thermophila* | XP_001020047.1 | MSNYLQKGFFIFVLLLVVNAKLGYDISVWEGEVNCFGCTKQSGYDFVVIEIYSSAKQFNNYLAQNIKSAYNASLEVDLYIFPDTTRDPAEQINSFFQYLIANETMSLFNNVWLDIENENLFFANCTQNIEFLQAMINTTENYISKDRIGLYASQHYWPIMCNTTVFSDLQLWYPRWDNQTNFNDFTPFGGFANPVMKQFYNDVQICCTNTDVDWRPDTQNNYFSNFSSRAFLKL |
| | XP_001025479.1 | MKFFAKLIILSVVASAVFCNFGIDIATLFNNYQCFVDNGYDLAIIRAYRSLGLVDANANTNLLNARAAGMDTDVYMFPCINDSKITAEQQVDDMINNLSLISSEDEGDAYDTIWVDVETNPSTTCSWNNISSVQDRCNYLQRIVNQILYRGKKVGIYASHYMWVQIFGDDKNCQNFNQLPLWYARYNNVTSTSDYSQYPFGGWTSAYAKQYAGSVSLCNVTVDLDYYN |
| *Tetratrichomonas gallinarum* | TEGa002098 | KVIDISAWQGDIDFKATKASGIRGVMIRSSYGTENPKQIDKQLANHATTAIAAGMLVGFYHYSYASNIQEAEAEADFCLKCIAPYKKDTVLPVAFDLEENSVAGLGKDYLAKMSIAFCRKIKAAGYKPMLYTNLNWARNYVNMAMIDAEGIDVWLAQYNSVCQYTGKHTMWQYTDHERIAGYGSGVDCSWCYVNYNDPDSPQMEQTQRGPNPSLVFNGKLDFYLTDLQYIYNLKQTAKADSTLYTKVKGETIKQGTSGQLTRWAQHRLSQLGFRCAESGTANSETMASIKQFQAYYQLGQGELGGTDWYYLLQP |

| | | |
|---|---|---|
| | TEGa014823 | ILDVSEHQDTINWTDAKEAGVKGVIIRAGFGSRNPSQIDKKIHNHALGALSANLAIGFYHYSY AKSVEIAKNETDFFFDIISEYKNRTTLPVAYDLEDKTIINWGLTKEDTATMAINFATKARSLGYI PMVYCNLNWAKNYINMKRVHEAGIELWIAQYNTHPQYIEGPIGMWQYSSKTHLPSYTEKFL DSSWCTRDYTNGTISFRENKQRGPNPKLEFRNNLDIDLGDIQSDLGVTKLRIANADSYYALT QYSVSLGTESFVVPFVQKRLKLLKFYDGDITGKADSATIQAINQFQKYYGLGQGTLQGTDW YYMIQP |
| *Tieghemostelium lacteum* | KYR00374.1 | MNRLLTLFALVVLLKVATATIGVDISSASDVPAFQCLKNEGYDFAIIRAWESVGQPDSNGPH SVYNAWEGGMADVDIYVFPCFSCGNGYGQVQSAIKYVQSYNAKFGMVWFDIEGPGTYWG GDQSSNAAFFADMVSGAQALGAHIGIYTSASQWIPIMGGYTGGSGFPLWYAHYDDNPSFS DFSPFGGWTAPAIKQYNGDLSVCGLGIDANWYP |
| *Trichomonas gallinae* | k32_R846165.p1 | LDVSYHNGTIDWNAVKQSDVKAVMIRSSYGKPDPSHVDIKFRENVLGALSAGLFIGIYHYTH ATTVSEAEAEAEFVLDIIKEFKDKITLPIAFDIEGEQREALSKDQNTKDAIAFCNKIREAGYTPM MYANLWFSGHHINMDEIRDANIDFWIAQWNSYTQYQEIFTMWQYSSTSKVPGCDGNIDSS WCSKDYTKGLIDFKVNKQKGLNPVYEFTKNSDTDVGELQSLLNLEKTWIPSELTYEEAAKIKI SPGESNSFVTWLQKRLTKLNFFNGSINGIFDSETETAVDKFHSFYELQPGIIS |
| | k32_S640890.p1 | TPFKIIDISHWQGDVDFAVAKAKSGIQGAMIRCGYGSPNPSQIDRQFKNHMREAIAANMLIGI YHYGYARTEAEAEAEAGFALSIIQDYKNYINFPVAYDVEDSTMIVGREQTTKNVLAFCRKIKA AGYKPMVYVNLLWTREYINMKAIDDAGIDVWFAEYNSYVKYTGNYTMWQFTDALSIAGYPE PVDASWVYKNYV* |
| | k48_S139806.p1 | MVPPPFKVIDISVWQGDIDFNAARVQSGIQGVMIRCGYGSYDPSQIDQQFHNHMRAAIACN MLIGIYHYGYARSPQEAENEADFCLSIIEPYRQYINLPVAYDVEDPTMNVGAWTVGQMAVAF GNKIRNAGYKPMLYCNLNWATNFIDMAQVNNAAIDVWLAQYNSECTYWGPYTMWQFTDA LWIGGYGSPVDASWCYVEYA* |
| | k48_S448125.p1 | PYKIIDVSHWQGDVNFTAAKEESEIQGAMIRCGFGRADPSHYDRQFQNHMREAIAARMLIGI YHYGYAQTVEQAEAEADFALSIIKDYRDDIIFPVAYDVEDSQMLVGKETITKMIIAFCRKIKDA GYKPMVYINYLWTQQYVDMAQIDAAGIDVWFAEYNSVPKYKGKFTMWQYTSSEKIAGYPD PVDA |
| | MRSU01007291.1 | PYKIIDVSHWQGDVNFTAAKEESEIQGAMIRCGFGRANPSHYDRQFQNHMREAIAARMLIGI YHYGYAQTVEQAEAEADFALSIIKDYKDDIVFPVAYDVEDSQMLVGKETITKMIIAFCRKIKDA GYKPMVYINYLWTQQYVDMAQIDAAGIDVWFAEYNSIPKYKGKFTMWQYTSSEKIAGYPDP VDASWVYYNY |
| | MRSU01003713.1 | PFKVIDISVWQGDIDFNAARVQSGIQGVMIRCGYGSYDPSQIDRQFHNHMRAAIACNMLIGI YHYGYARSPQEAENEADFCLSIIEPYRQYINLPVAYDVEDPTMNVGAWTVGQMAVAFGNKI RNAGYKPMLYCNLNWATNYIDMSQVNNAAIDVWLAQYNSQCTYWGPYTMWQFTDALWIG GYGSPVDASWCYVEY |
| | MRSU01001113.1 | MAYKVVDVSVWQGKIDFNAAKTYSGIRGVMIRSGFGSKAGQEDKQFINHLREAIAANIAVGI YHYGYARTEAQAIAEADFCHNIINPYKGYISMPVAYDVEDATMNVGRAQVARNAIAFCNRM |

| | | |
|---|---|---|
| | | RAYGYKPMIYCNKNWATNYIDMSQINNAGIDVWIAQYNSVCTYTGPYVMWQFTNSLRISGY SSGVDGSYAYK |
| *Trichomonas stableri* | WGY81450.1 | MPYKVIDISVWQGDIDFNAARDQSGVQGVMIRCGYGSYSPSQIDGQFHNHMRAAIAANMLI GVYHYGYARSVAEAEGEADFAISIFDQYRQWIRLPIAYDIEDPTMNVGRDLVGQMAVAFAN KIRAAGYLPMCYCNLNWATNFINMDIVNANGVDVWMAQYNSQCTYWGPYVMWQFTDALP IGGYYAGVDGSWCYKDY |
| | WGY81646.1 | MNAYKIIDVSVWQGAIDFNAAKVNSGIQGFMIRAGFGSAAGQEDKQFANHMREAIAANILIG AYHYGYARTEAQAIAEADFCAKIINPYKGYFRMPVAYDVEDATMNVGKVQTAKNAIAFCNR MKAYGYKPMIYCNLNWARNYIDMSMIDAAGIEVWIAQYNSVCQYTGPHTMWQYTNALRIS GYSSGVDGSWVYKNY |
| | WGY81674.1 | MRDLPFKIIDVSKWQREIDFQAAKDNSQIQGVMIRSSYGSYDPKQIDTMFANHMRNAIAANM LIGTYHYGYARTVEQAENEADFAISVIKPYKDYIKFPVAYDVEDAAMNVGKELVADMVIAFCN KVKAAGFKPMLYINLQWINNYINIDRINEAGIDVWFAEYNSYCKYTGPKTMWQFTDALSISG YSEPVDASWVYKNYE |
| *Trichomonas tenax* | k32_J419651.p1 | PYKVVDVSVWQGNIDFNAAHINSGINGVMIRAGYGSHVGQEDKQFVNHLREAIAAKIAIGAY HYGYAKTEAQAIAEADYCHSIINPYKAHINMPVAYDVEDATMAVGKVQVAKNVLAFCNRMK AYGYKPMLYCNLNWATNYIDMTMVNNAGIEVWIAQYNSVCTYKGPYVMWQFTNALRISGY TSGVDGSYAYKNW* |
| | k32_R410566.p1 | MVAPPFKVIDISVWQGDIDFNAARAESGIQGVMIRCGYGSYDPSQIDGQFHNHIRAAIACNM LIGIYHYGYASSPQEAEDEADFCLSIIEPYRQWINLPVAYDVEDPTMNVGAWTVGAMAVAFG NKIRNAGYKPMLYCNLNWATNFIDMDQVNNAAIDVWMAQYNTECTYWGPYTMWQFTDAL WIGGYSSPVDASWCYVEYA* |
| | k48_S124239.p1 | RHAYIEACHFSILMPATPYKIIDVSHWQGDINFSLAKDKSEIQGAMIRCGFGTENPAHYDRQF QNHMREAIAAHMLIGIYHYGYARTVEEAEAEADFALKIIEDYRPNIVFPVAYDVEDAQMMVG KETITKMILAFCKKIKDAGYKPMVYVNLLWTREYIDMAQIDAAGIDVWFAEYNTYPKYQGKFT MWQFTNHQEIAGYPDPVDASWVYYNYEA* |
| | k64_S51806.p1 | MAFRVIDVSKYQPSLNWNGIKSSGVKAVMVRTSYGKRDPSHVDPMFRNHTIGALTAGLFV GFYHYTHATTVSEAESEADFVIDIIKEFKKNTTFPVAFDLEGSSRESLPRNQNTENTIAFCNKI RNAGYTPMLYCNLNFATNFIYMDQIRSANIDYWNAQYYSRPKYQNQFTMWQYSSQTRVPG YDGDIDSSWCSKDYTNGLINFKVNSQRGIDPALTFTKNSDTDVGELQSLLGIDKTWIPSYLT YQTAVQKEIRLGDKSDLVEWLQSRLMKLKFYNGQVNGYFDAETEQAVNSFHSYYELQHGC ISGIDWYYLIQP* |
| | JALKBU010000432.1 | PFKLIDVSHWESDIDFQAAKDQSGIQGVMMRCGYGSPNPAVDRMFVNHMREAIAARMLIGI YHYGYARTEEQAEAEANFALSIIKDYKDNIIFPVAYDVEDAQMIVDRDQVTKNVLAFCRKIKE AGYKPMLYINLLWTREYINMDQIDAAGIDVWFAEYNSYVKYTGNYTMWQFTDGLPIAGYPE PVDASWVYKNY |

| | | |
|---|---|---|
| *Trichomona vaginalis* | TVAGG3_0398000<br>TVAG_088640 | MPYKVIDISVWQGDIDFNAARDQSGVQGVMIRCGYGSYSPAQIDGQFHNHMRAAIAANMLI<br>GVYHYGYAQSVPEAEGEADFAISIFDQYRQWIRLPIAYDIEDPTMNVGRDLVGQMAAAFAN<br>KIRAAGYTPMCYCNLNWATNFINMDIVNANGIDVWIAQYNSECQYWGPYVMWQFTDCLPI<br>GGYYAGVDGSWCYKDY |
| | TVAGG3_0440180 | MNAYKIIDVSVWQGNIDFNAAKVNSGIQGVMIRAGFGSAAGQEDKQFANHMREAIAANILIG<br>AYHYGYARTEAQAIAEADFCAKIINPYKSHFRMPVAYDDEDATMNVGKVQTAKNAIAFCNR<br>MKAYGYKPMVYCNLNWARNYIDMSMINAAGIEVWIAQYNSVCQYTGPHTMWQYTNALKIS<br>GYSACVDGSWVYKNY |
| | TVAGG3_0441060 | MPDLPFKIIDVSKWQNEIDFQAAKDNSQIQGVMIRSSYGSYDPKQIDTMFENHMRNAIASNM<br>LIGTYHYGYARSVEQAEREADFAISVISPYKDYIKFPVAYGVEDAAMNVGKELVADMVIAFC<br>NKVKAAGFKPMLYLNLVWANNYCNMDRINEAGIDVWLAEYNSYCKYTGPKTMWQFTDALS<br>ISGYPEPVDASWVYKNYE |
| | TVAGG3_1010370 | MVATPFKVIDISVWQGDIDFNAARDQSGIQGVMIRCGYGSADPSQVDKQFVNHIRAAIACN<br>MLIGIYHYGYAENVQEAENEADFALSIIEPYKQYITLPVAYDIEDPTMNVGRDLVGQMAVAFA<br>NRIRAAGYRAMTYCNLNWATNYINMDWINNAGIDVWMAQYNSQCTYWGPYTMWQFTDA<br>QPIGGYYAGVDCSWCYVAYA |

**Table S2.** List of primers.

| Experiment | Gene | Accession no. | Vector | Note | Primer forward | Primer reverse |
|---|---|---|---|---|---|---|
| Viability assay | GH25 | TVAG_088640 | pTagVag-V5-Pur | | 5'-TACGCATATGCCGTACAAGGTTATCGACAT-3' | 5'-GCATGGATCCGTAATCCTTATAGCACCAA-3' |
| | HCP1 | TVAG_206500 | pTagVag-HA-Neo | | 5'-TACGCATATGCTTTCCTCCCAACCGGCAT-3' | 5'-GCATGGATCCCTGAAGGATATCCTTGAGATCATCCTC-3' |
| qPCR | GH25 | TVAG_088640 | | | 5'-GCCGATTTCGCTATCAGTATTT-3' | 5'-TGACCTACCAAATCTCTTCCTAC-3' |
| | TBSR1 | TVAG_166850 | | | 5'-ACTATGGCCACCACGTTATTA-3' | 5'-GAGCTTGGGAAGGTTTCATTT-3' |
| | TBSR12 | TVAG_369130 | | | 5'-CCCTTTGGTGGAAAGAGATAAC-3' | 5'-TCGGTGTTCGTGAAGAATTTAG-3' |
| Mutagenesis PCR for mutTvGH25 preparation | GH25 | TVAG_088640 | | Fragment 1 | 5'-TACGCATATGCCGTACAAGGTTATCGACAT-3' | 5'-TTGTTGGATCTTGGATAGCGTAA-3' |
| | | | | Fragment 2 | 5'-TTACGCTATCCAAGATCCAACAA-3' | 5'-GCATGGATCCGTAATCCTTATAGCACCAA-3' |

**Dataset S1 (separate file).** Proteins identified by mass spectrometry analysis upon co-incubation of *T. vaginalis* with *L. jensenii*.

**Dataset S2 (separate file).** Proteins identified by mass spectrometry analysis in the control (*T. vaginalis* without *L. jensenii)*.

### SI References


1. B. D. Janssen, *et al.*, CRISPR/Cas9-mediated gene modification and gene knock out in the human-infective parasite *Trichomonas vaginalis*. *Sci. Rep.* **8** (2018).

2. J. Schindelin, *et al.*, Fiji: An open-source platform for biological-image analysis. *Nat. Methods* **9**, 676–682 (2012).

3. J. Cox, *et al.*, Accurate proteome-wide label-free quantification by delayed normalization and maximal peptide ratio extraction, termed MaxLFQ. *Mol. Cell. Proteomics* **13**, 2513–2526 (2014).

4. J. Alvarez-Jarreta, *et al.*, VEuPathDB: the eukaryotic pathogen, vector and host bioinformatics resource center in 2023. *Nucleic Acids Res.* **52**, D808–D816 (2024).

5. J. Štáfková, *et al.*, Dynamic secretome of *Trichomonas vaginalis*: Case study of β-amylases. *Mol. Cell. Proteomics* **17**, 304–320 (2018).

6. R. Kolde, pheatmap: Pretty Heatmaps. (2018).

7. K. J. Livak, T. D. Schmittgen, Analysis of Relative Gene Expression Data Using Real-Time Quantitative PCR and the 2−ΔΔCT Method. Methods 25, 402–408 (2001).

8. M. Varadi, *et al.*, AlphaFold Protein Structure Database: massively expanding the structural coverage of protein-sequence space with high-accuracy models. *Nucleic Acids Res.* **50**, D439–D444 (2022).

9. J. Jumper, *et al.*, Highly accurate protein structure prediction with AlphaFold. *Nature* **596**, 583–589 (2021).

10. A. M. Lau, *et al.*, Exploring structural diversity across the protein universe with The Encyclopedia of Domains. *Science* **386**, eadq4946 (2024).
